# The systemically induced sugar transporter SWEET11 regulates growth-defense trade-offs during *Serendipita indica* symbiosis in Arabidopsis

**DOI:** 10.64898/2026.07.21.739764

**Authors:** Abhimanyu Jogawat, Sidhardh H. Menon, Madhan Sanyasi, Divya Goyal, Athira Mohandas Nair, Jyothilakshmi Vadassery

**Affiliations:** BRIC-National Institute of Plant Genome Research, New Delhi-110067

## Abstract

Sugar exchange at the root interface is a pivotal process governing the establishment and stability of plant-fungal symbioses. Precise regulation of sugar exchange determines the success of this ecologically significant interaction. Sugar Will Eventually be Exported proteins (SWEETs) constitute a family of regulatory, energy-independent bidirectional sugar transporters that influence plant development, stress resilience, and survival. However, how specific SWEET transporters coordinate systemic carbon allocation and immune regulation during beneficial plant-fungal interactions remains poorly understood. In this study, we examined the role of the systemically induced Arabidopsis sugar transporter SWEET11 during association with the beneficial endophytic fungus *Serendipita indica* and following treatment with its elicitor, cellotriose (CT). Expression profiling of SWEET family members revealed a rapid and preferential induction of SWEET11 in aerial tissues upon fungal colonization and CT treatment. Loss-of-function of SWEET11 compromises key mutualistic outcomes, including plant growth enhancement, fungal colonization efficiency, penetration ability, carbohydrate distribution, and the regulation of defense-related phytohormones such as jasmonic acid and abscisic acid. Global transcriptome analysis further demonstrated that SWEET11 regulates whole-plant responses by orchestrating genes involved in central metabolism, secondary metabolite production, sesquiterpenoid and triterpenoid pathways, as well as defense signaling and nutrient transport systems. We show that SWEET11 interacts with a stress associated SNF1-related protein kinase (SnRK2.8) and plays a crucial role in enabling fungal establishment while mitigating host defense responses, and supporting plant growth. Our data shows that SWEET11 functions as a shoot-derived sugar exporter that directs carbon toward roots, facilitating sugar unloading to *S. indica*. This controlled carbon supply allows the fungus to meet its metabolic demands without disrupting host sugar balance, thereby maintaining a stable and well-regulated symbiotic association under immune constraints.

## 1. Introduction

*Serendipita indica* (syn. *Piriformospora indica*), a growth-promoting fungus native to the Thar Desert, functions as a generalist root endo-mutualist with broad host compatibility (Verma et al. 1998; Kundu and Vadassery 2022). Interactions of *S. indica* with diverse host plants have been extensively studied in the context of locally induced defense responses in roots as well as at the whole-plant level. During the symbiotic phase, *S. indica* actively modulates a wide range of host cellular and metabolic processes. These include calcium signaling, glucosinolate accumulation, callose deposition, reactive oxygen species (ROS) homeostasis, ion and sugar balance, phytohormone biosynthesis and signaling, and the production of secondary metabolites, all of which contribute to successful colonization and accommodation within plant roots (Vadassery et al. 2008; Jacobs et al. 2011; Johnson et al. 2018; Jogawat et al. 2020; Opitz et al. 2021; Ortega-Villaizán et al. 2026). *S. indica* exerts profound effects on distal plant tissues via systemic signaling and promotes systemic acquired resistance (SAR) through the accumulation of defense-related metabolites, modulation of phytohormone pathways, and activation of antioxidant systems (Waller et al. 2008; Pedrotti et al. 2013; Vahabi et al. 2015; Li et al. 2022). It influences salicylic acid signaling by modulating Nonexpressor of Pathogenesis-Related genes 1 (NPR1) localization, thereby contributing to induced resistance in aerial tissues (Stein et al. 2008; Zhang et al. 2024). In addition to modulating immune signaling, *S. indica* also reprograms host primary metabolism, particularly sugar metabolism, by altering sugar levels and increasing their availability in the apoplast (Strehmel et al. 2016; Opitz et al. 2021; Kundu et al. 2022; Jogawat et al. 2026). Apoplastic sugar dynamics are increasingly recognized as important components of plant defense against biotic stress (Yamada and Mine, 2024; Mao et al. 2025). Plants possess a diverse repertoire of sugar transporters that mediate the mobilization of sucrose, mono-, and disaccharides. Beyond their essential roles in growth and development, these transporters contribute significantly to stress adaptation and defense responses (Doidy et al. 2012; Julius et al. 2017; Yamada and Mine, 2024). Importantly, plant-associated microorganisms often deploy effectors to manipulate host sugar transport, thereby increasing sugar availability in the apoplast to support their colonization. Among sugar transporter families, Sugar Will Eventually be Exported transporters (SWEET) have been extensively studied for their roles in pathogenesis, as well as for being targets of fungal and bacterial effectors. While SWEET proteins are primarily involved in physiological processes such as nectar secretion, pollen development, pollen-tube growth, and seed filling, they also play a dual role in plant-pathogen interactions and abiotic stress tolerance (Chen et al. 2010; Baker et al. 2012; Eom et al. 2015; Chen et al. 2022). Importantly, SWEET11, SWEET12 and SUC2 facilitates sucrose efflux from phloem (Chen et al. 2012). SWEET11 and SWEET12 are localized to parenchymal cell membrane whereas SUC2 is on the companion cell membrane of phloem (Chen et al. 2012; Stadler and Sauer, 1996). These transporters are crucial for efficient phloem loading and long-distance transport (Chen et al. 2012; Stadler and Sauer, 1996). Plant-associated microorganisms often exploit host sugar transport systems to redirect carbon flow toward infection sites, thereby supporting their survival, proliferation, and successful invasion (Walerowski et al. 2018; Fatima and Senthil-Kumar 2021). To achieve this, many microbes secrete effector molecules that specifically target and modulate SWEET transporters as part of their infection strategy to access host-derived sugars (Streubel et al. 2013). In rice, *Os*SWEETs are well characterized targets of the transcription activator-like (TAL) effectors, including PthXo1, PthXo3, TalC, Tal5, and AvrXa7 secreted by *Xanthomonas oryzae*. These effectors mainly induce the expression of *OsSWEET11* (*Xa13*) and *OsSWEET14* (*Os11N3*) thereby enhancing sugar efflux to facilitate pathogen nutrition and virulence (Chu et al. 2006; Yang et al. 2006; Antony et al. 2010; Bogdanove et al. 2010; Talbot 2010; Verdier et al. 2012; Streubel et al. 2013). Consistently, disruption of *Os*SWEET11 has been shown to confer resistance to sheath blight in rice (Ponnurangam et al. 2026).

The involvement of SWEET transporters in symbiotic interactions has been well documented, particularly in AM and rhizobial symbioses (Desrut et al. 2020; Loo et al. 2024; Zhu et al. 2026). Multiple SWEET members have been identified as key regulators of sugar allocation to symbiotic partners across diverse plant species. For example, *Gm*SWEET3c and *Gm*SWEET38 in soybean, *Mt*SWEET1b in *Medicago truncatula*, *Lj*SWEET3 in *Lotus japonicus*, *St*SWEET7a in potato, and *Do*SWEET14 in *Dendrobium officinale* have been shown to play critical roles in carbohydrate partitioning during symbiosis (An et al. 2019; Tamayo et al. 2022; Zhu et al. 2026; Chen et al. 2025; Li et al. 2025). In soybean roots, the *At*SWEET11 homolog, *Gm*SWEET6 facilitates sugar export to AM fungi (Zheng et al. 2024), while in *M. truncatula*, *Mt*SWEET11 localizes to the nodule vasculature, supporting its role in symbiotic nitrogen fixation (Kryvoruchko et al. 2016). Moreover, *At*SWEET11 has been demonstrated to be important for non-obligate symbiosis in *Arabidopsis* (Desrut et al. 2020). Although root transcriptomic studies have reported the induction of *SWEET11* during *S. indica* colonization, its functional role in this symbiosis remains unclear (Pérez-Alonso et al. 2022; Gandhi et al. 2024; Jogawat et al. 2026), and its precise contribution to sugar allocation and systemic signaling remains uncharacterized.

In the present study, we investigated whether root colonization by *S. indica* triggers systemic reprogramming of sugar transport and defense signaling in the shoot to support mutualistic growth promotion. We identified SWEET11 as a key sugar transporter that is systemically induced in aerial tissues in response to root colonization by *S. indica*, along with SnRK2.8. Our findings uncover a systemic SWEET11–ABA–SnRK2.8 regulatory module that integrates sugar allocation and signaling to balance growth and defense, thereby ensuring the successful establishment and maintenance of *S. indica* symbiosis.

## 2. Materials and methods

### 2.1. Fungal strain, Plant and Growth Conditions

We utilized *Serendipita indica* (syn. *Piriformospora indica*; Verma et al. 1998) which was maintained and cultivated on modified Kaefer’s medium (Hill and Kafer 2001) at 28°C with constant shaking of 110 rounds per minutes. For co-cultivation and growth promotion experiments, *Arabidopsis thaliana* (Columbia ecotype; Col-0; wild type) and its mutant lines were used. We used T-DNA insertion lines of *SWEET11* (At3g48740; SALK_073269.20.35.x; Chen et al. 2012) and confirmed *SWEET11* silencing by RT-PCR (**Fig. S1**). This mutant line has previously been utilized in other studies (Desrut et al. 2020). Additionally, the confirmed T-DNA insertion lines of *SnRK2.8* (At1g78290) i.e. *snrk2.8-1* (SALK_069354C; Shin et al. 2007) and *snrk2.8-2* (SALK_073395C; Shin et al. 2007) and *CORK1* (At1g56145) i.e. *cork1* (SALK_021490C; Tseng et al. 2022) were obtained from Arabidopsis Biological Resource Centre (https://abrc.osu.edu/). For growth assay, the co-cultivation of Arabidopsis with *S. indica* was performed on 1X PNM solid media plates. For this, the surface-sterilized seeds first stratified and then germinated on half-strength MS medium for 7 days before co-cultivation. *S. indica* co-cultivation was performed as per procedure mentioned earlier (Johnson et al. 2011; Jogawat et al. 2020, 2026). The samples from different conditions were collected at different time points such as 2, 7, 14, 21, and 30 days post inoculation (dpi) as per experimental needs.

### 2.2. Expression of SWEETs upon CT- and *S. indica*-treatment in systemic and local tissues

We added cellotriose (CT) in liquid ½-strength MS medium for a final 10 µM concentration and used the solution to treat ten days old seedlings. The samples were collected at 0 min, 30 min, 1 hr, 2 hr, 4 hr, and 8 hrs. Shoots were harvested at 7 and 14 dpi and immediately frozen in liquid N_2_. Total RNA was extracted and cDNA was prepared as mentioned earlier (Jogawat et al. 2026). Further, iTaq Universal SYBR Green Mix (Bio-Rad) with gene-specific primers (**Supplementary Table S1**) was used in a 96-well plate on a CFX96 RT-PCR Detection System (Bio-Rad). For transcripts normalization, *AtActin2* (*At3g18780*) primers pair was used as endogenous control and the relative expression of genes was calculated (Livak and Schmittgen, 2001).

The roots of *SWEET11* promoter with β-glucuronidase (GUS) fusion transgenic line (Le Hir et al. 2015) were treated with 10 µM CT supplemented in half-strength MS medium. These seedlings were also co-cultivated with *S. indica*. At 2 dpi and 7 dpi, they were harvested, vacuum-infiltrated with GUS staining solution (Li 2011), and incubated in the dark at 37°C for 12 hours. The tissues were treated with a freshly prepared decolorizing solution [ethanol: 50% glycerol: acetone (9:3:3)] at room temperature and finally observed under a Stereo zoom Microscope (Nikon).

### 2.3. Estimation and detection of *S. indica* colonization

*S. indica*-colonized roots were carefully harvested and washed. Alexa Flour 488 WGA fluorescent dye was used to stain fungal spores. Finally, the roots were mounted on glass slides to observe under Laser Scanning Microscope (AOBS TCS-SP8; LEICA GERMANY) using emission at 505–530 nm and excitation at 470 nm with optical sectioning at 1µm per step across the root. For penetration assay, 3D images were generated by staking optically sectioned images into z-plane with scale of each image representing the depth penetrated by *S. indica*. Using a Real time PCR-based approach, we also performed relative fungal load estimation as previously described (Jogawat et al. 2020; Kundu et al. 2022; Jogawat et al. 2026). In brief, *Arabidopsis Actin2* (*At3g18780*) and *S. indica Tef1* primers were used in RT PCR. By using CT value of *S. indica*-specific gene *SiTef1*, the fold changes in relative fungal DNA amount were calculated by normalizing CT value of *SiTef1* against CT value of *Arabidopsis Actin2* with the ΔΔCT equation (Bütehorn et al. 2000; Vadassery et al. 2008; Jogawat et al. 2020; 2026).

### 2.4. Soluble sugars levels analysis by targeted GC-MS

The sugar estimation was done using GC-MS/MS, according to previously mentioned procedure, in both shoot and whole seedlings (Schauer et al. 2005; Kundu et al. 2022; Jogawat et al. 2026).

### 2.5. Phytohormone estimation

Defense phytohormones such as ABA, JA, JA-Ile and SA were estimated as mentioned in our earlier studies (Vadassery et al. 2012; Jogawat et al. 2026).

### 2.6. Yeast two-hybrid assay

The coding sequences corresponding to the C-terminal regions of SWEET11 and SWEET12 were individually inserted into the pGBKT7-BD vector (Clontech Laboratories Inc.) to generate fusions with the GAL4 DNA-binding domain. The coding sequences of SnRK2.8 and the kinase domain of CORK1 (CORK1-KD) were independently cloned into the pGADT7-AD vector (Clontech Laboratories Inc.), placing them downstream of the GAL4 activation domain. All recombinant plasmids were generated by restriction enzyme-based cloning using *EcoRI* and *BamHI* restriction sites, followed by ligation with Phusion Clonase (Thermo Fisher Scientific, USA). Primer sequences used for amplification are provided in **Supplementary Table S1**. The bait and prey plasmids were co-transformed into the Y2HGold yeast strain using the Matchmaker Gold Yeast Two-Hybrid System (Clontech Laboratories Inc.). Transformants were initially selected on double-dropout (DDO) medium lacking leucine and tryptophan (-Leu/-Trp), and successful transformation was verified by colony PCR with gene-specific primers (Supplementary Table S1). As experimental controls, the positive control consisted of co-transformation of pGBKT7-T antigen with pGADT7-p53, whereas pGBKT7-T antigen together with pGADT7-Lam served as the negative control. Additional negative controls included yeast cells transformed with the corresponding empty pGBKT7-BD and pGADT7-AD vectors. Confirmed transformants were cultured in liquid DDO (-Leu/-Trp) medium at 30°C for 48 h. The cultures were normalized to an OD□□□ of 2.0 and serially diluted to OD□□□ values of 1, 0.1, 0.01, 0.001, and 0.0001. Aliquots (10 μL) of each dilution were spotted onto DDO (-Leu/-Trp) and triple-dropout (TDO; -Leu/-Trp/-His) selective media. The plates were incubated at 30°C for 3 days before assessing yeast growth. Each yeast two-hybrid experiment was independently repeated at least three times to ensure reproducibility.

### 2.7. Split Luciferase assay

For the luciferase complementation imaging (LCI) assay, the full-length coding sequences of SWEET11 and SnRK2.8 were cloned into the pCAMBIA-nLUC and pCAMBIA-cLUC vectors, respectively. The primer sequences used for cloning are provided in **Supplementary Table S1**. Recombinant plasmids were introduced into *Agrobacterium tumefaciens* strain GV3101 by the heat-shock transformation method. Transformed bacterial cultures were pelleted by centrifugation at 2200 × g for 15 min and resuspended in infiltration buffer containing 1 M MgCl□, 100 mM acetosyringone, and 1 M MES-KOH (pH 5.7). The suspensions were incubated in the dark at 22 °C for 3 h before infiltration. Agroinfiltration was performed on the abaxial side of leaves from 6-week-old *Nicotiana benthamiana* plants. At 2 days post-infiltration, the leaves were infiltrated with 1 mM luciferin and incubated in the dark for 7 min. Luciferase signals were detected using a Bio-Rad low-light cooled CCD imaging system, and signal intensities were quantified with ImageJ software. The assay was repeated in three independent biological experiments, each comprising 6–8 plants.

### 2.8. BiFC assay

Bimolecular fluorescent complementation (BiFC) assay was performed as mentioned previously (Xu et al. 2016; Meena et al. 2019). Briefly, to generate BiFC constructs for transient co-expression, the full-length CDS of *SWEET11* and *SnRK2.8* was cloned into pSITE-nEYFP-C1 / CD3-1648 and pSITE-cEYFP-C1 / CD3-1649 to obtain *SWEET11*-nEYFP and *SnRK2.8*-cEYFP, respectively. The constructs, along with p19, were transformed into *Agrobacterium tumefaciens* strain GV3101. Agrobacterium transformants with *AtSWEET11*, and *SnRK2.8* were used to infiltrate *Nicotiana* plant leaves. Additionally, we used negative (CNGC19-CML42) and positive (CNGC19-CAM2) interactions. The infiltrated area was imaged after 48 hrs using a confocal laser scanning microscope AOBS TCS-SP8 (LEICA GERMANY).

### 2.9. Total RNA isolation, library preparation and RNAseq analysis

To analyse differentially expressed genes (DEGs) upon *S. indica* colonization with Col-0 and *sweet11*, the colonized roots and non-colonized control roots were pooled (each sample up to 250 mg) separately and total RNA were isolated as mentioned previously (Jogawat et al. 2020). After quantity, integrity and quality assessment of RNA, they were processed for cDNA library (PE150) construction and RNA sequencing on Illumina NovaSeq™ 6000 platform. The processing of the obtained raw fastq reads was performed as mentioned in our previous study (Jogawat et al., 2026 and references therein). The annotated DESeq2 result file was filtered based on two parameters such adjusted *p*-value of False Discovery Rate (FDR) at ≤ 0.05 and Log2FoldChange (Log2FC) at ± 0.58. The significant DEGs from the groups were subjected to gene ontology (GO) and ‘Kyoto Encyclopedia of Genes and Genomes’ (KEGG) enrichment analysis with the *Arabidopsis thaliana* genome, a significance threshold of 0.05, and FDR as the adjustment method.

### 2.10. Bioinformatics analysis of transcriptome data

For KEGG and GO analysis, we used online bioinformatics tools such as ShinyGO (Ge et al. 2020, bioinformatics.sdstate.edu/go/) and AgriGO (systemsbiology.cau.edu.cn/agriGOv2/). For clustering of DEGs of all conditions, we used iDEP online tool (bioinformatics.sdstate.edu/idep/). For Heat maps and Venn diagrams generation, we utilized TBtool (Chen et al. 2020) and DiVenn 2.0 tool (https://divenn.tch.harvard.edu/v2/). We also predicted subcellular localization of the DEGs utilizing SUBA5 tool (https://suba.live/).

### 2.11. Statistical analysis

GraphPad Prism version 8.4.2 software tool used for the graphs and plots generation. For unpaired or paired two-tailed Student’s *t*-test, Microsoft’s excel sheet was used. For ANOVA analysis GraphPad software was used for calculating statistical significance (*p*-value ≤0.05) between or among control and treated samples. Asterisks or different letters were put to indicate statistical significance difference in graphs and plots.

## 3. Results

### 3.1. *S. indica* colonization systemically induces SWEET expression in shoots

Sugars are synthesized in shoot during photosynthesis and transported to roots (Hennion et al. 2019). To identify systemically induced sugar transporter genes involved in the symbiosis between *S. indica* and *Arabidopsis thaliana*, we analyzed *SWEET* gene expression in the shoots of colonized and non-colonized plants at 7 and 14 days post-inoculation (dpi). For this, we harvested shoots and checked systemic induction of all *SWEET* genes. Among all genes analyzed, we observed the induction of seven *AtSWEETs* which include *SWEET4*, *SWEET8*, *SWEET11*, *SWEET12*, *SWEET13*, *SWEET14*, and *SWEET15* in systemic shoots in response to *S. indica* root colonization. Notably, the expression of *SWEET11* (4.18-folds) was the highest among the induced *SWEETs*, along with *SWEET4* (3.82-folds) and *SWEET13* (3.97-folds) in *S. indica*-colonized plants’ shoot at 7 dpi (**Fig. 1A**). At 14 dpi, SWEET6, SWEET9, SWEET11, and SWEET15 were found to be induced. The expression profiling indicates the possible involvement of these SWEET transporters *S. indica*-mediated sugar mobilization and suggests *SWEET11* is an important sugar transporter for further study in the context of *S. indica* colonization, due its proven role in phloem loading (Chen et al. 2012).

**Fig. 1.**
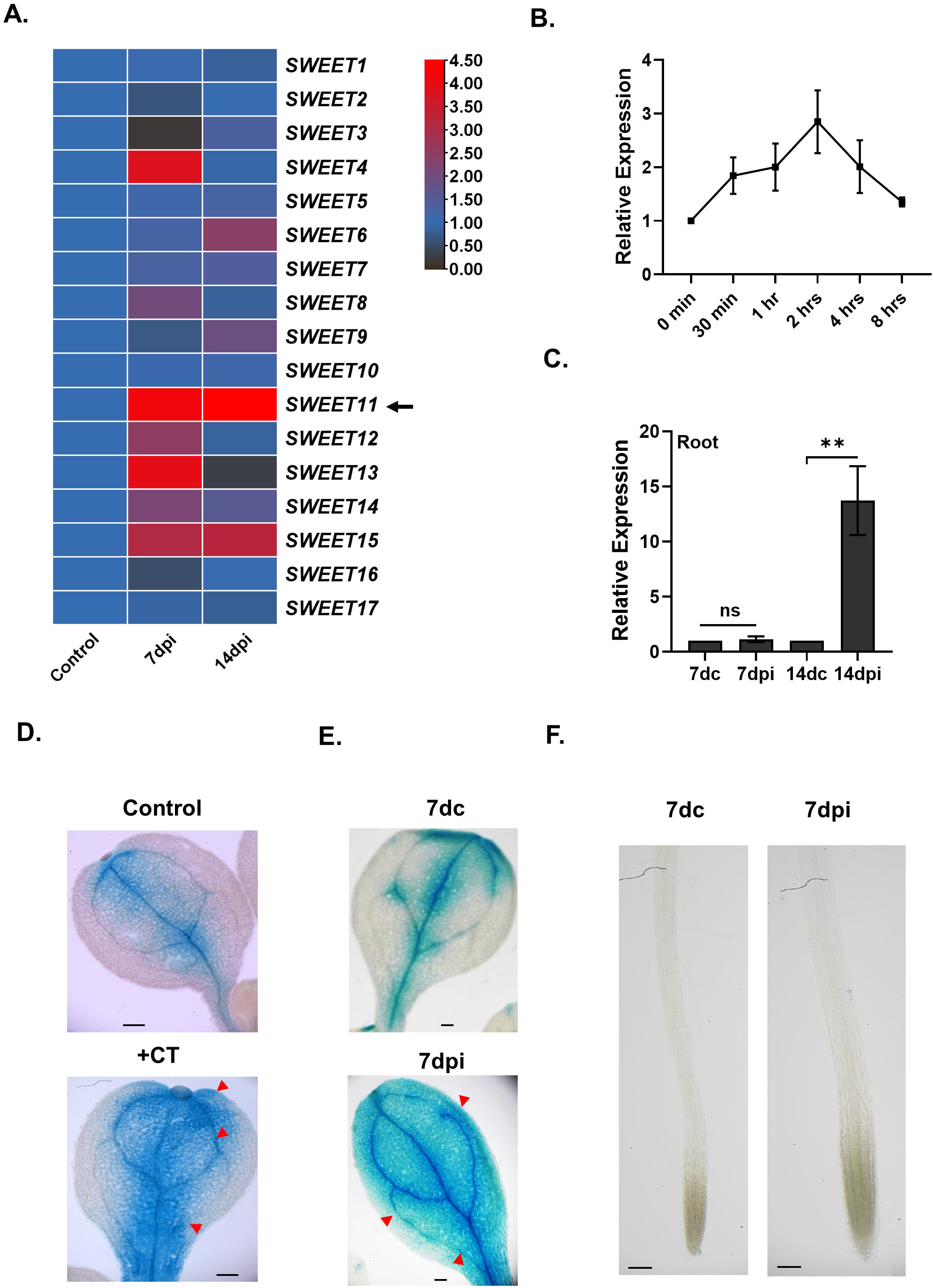
Expression analysis of *SWEET*s upon *S. indica* colonization: **A.** Seven days old seedlings were exposed to *S. indica* plugs and the relative expression of *SWEET* genes was estimated in shoots at 7 dpi. Data represents mean fold change ± SEM (N = 3-4, 4*10). **B. Time course expression analysis of *SWEET11* post CT-treatment:** Ten days old seedlings were exposed to 10 µM CT and *SWEET11* expression was checked at 0, 30 min, 1 hr, 2 hr, 4 hr and 8 hrs post CT-treatment. Data represents mean fold change ± SEM (N = 3-4, 4*10). **C. Expression of *SWEET11* in colonized roots.** Root of *S. indica*-colonized seedlings were harvested and the relative expression of *SWEET11* was estimated at 7 and 14 days post inoculation (dpi) compared to their respective controls at 7 and 14 days controls (7 dc and 14 dc) post inoculation. Data represents mean fold change ± SEM (n = 4, 4*10). **D. *SWEET11* promoter activity upon CT-treatment:** 10 days of seedlings expressing p*SWEET11*-GUS were treated with 10 μM CT for 2 hrs and GUS assay was performed. Blue color and arrow indicates induced GUS expression in different tissues. **E, F. *SWEET11* promoter activity upon *S. indica* colonization.** Seedling were grown for 7 days on solid ½ MS and then co-cultivated with *S. indica*. *SWEET11* promoter activity at 7 dpi in leaf (**E**) and root tip (**F**). Blue color and arrow indicates GUS expression (Scale bar = 100 μm).

### 3.2. Both cellotriose and *S. indica* treatment induces *SWEET11* predominantly in shoot

To test whether *SWEET11* is induced by *S. indica*-originated elicitor cellotriose (CT), we applied CT (10 μM) and checked *SWEET11* expression at different time points. *SWEET11* transcript levels peaked at 2 h after CT treatment and declined thereafter (**Fig. 1B**). Previous studies have also reported *SWEET11* induction in roots (Pérez-Alonso et al. 2022; Gandhi et al. 2024; Jogawat et al. 2026). Consistent with this, SWEET11 expression peaked at 14 dpi in roots (**Fig. 1C**), whereas in shoots it peaked earlier, at 7 dpi (**Fig. 1A**). Further, to investigate the tissue-specific promoter activity of *SWEET11*, *ProSWEET11*:*GUS* line (Le Hir et al. 2015) was used. Following the CT treatment, *SWEET11* promoter activity was predominantly observed in the shoot, particularly in leaf veins and petioles (**Fig. 1D**). Likewise, during *S. indica* colonization, *SWEET11* promoter activity was strongly induced in the shoot, with very little induction in the roots at 2, 7 and 14 dpi (**Fig. S2A-B, Fig. 1E-F and Fig. S3A-B**). These findings suggest that *S. indica* systemically induces *SWEET11* expression in a tissue-specific manner, with *SWEET11* primarily functioning in the shoot.

### 3.3. *S. indica* fails to promote growth in *sweet11* mutants

SWEETs are involved in sugar transport during mycorrhizal symbiosis (An et al. 2019; Tamayo et al. 2022; Chen et al. 2025). To investigate functional relevance of *SWEET11* in *S. indica*-mediated growth promotion, *sweet11* mutant plants were co-cultivated with *S. indica* and growth was assessed at 14 dpi. While *S. indica* significantly enhances growth in WT plants, no growth promotion was observed in the *sweet11* mutants, compared to WT control (**Fig. 2A-B**). Both shoot and roots weights of *sweet11* mutants do not exhibit any enhancement on *S. indica* colonization, compared to that in WT (**Fig. S4A-B**). This observation was further validated in soil-based co-cultivation experiment at 30 dpi, which also reiterated that *sweet11* mutant plants have no growth promotion upon *S. indica*-colonization, in contrast to the pronounced growth enhancement seen in WT plants (**Fig. S5A-B**). This underscores the critical role of *SWEET11* in mediating *S. indica*-induced growth benefits to host plant.

**Fig. 2.**
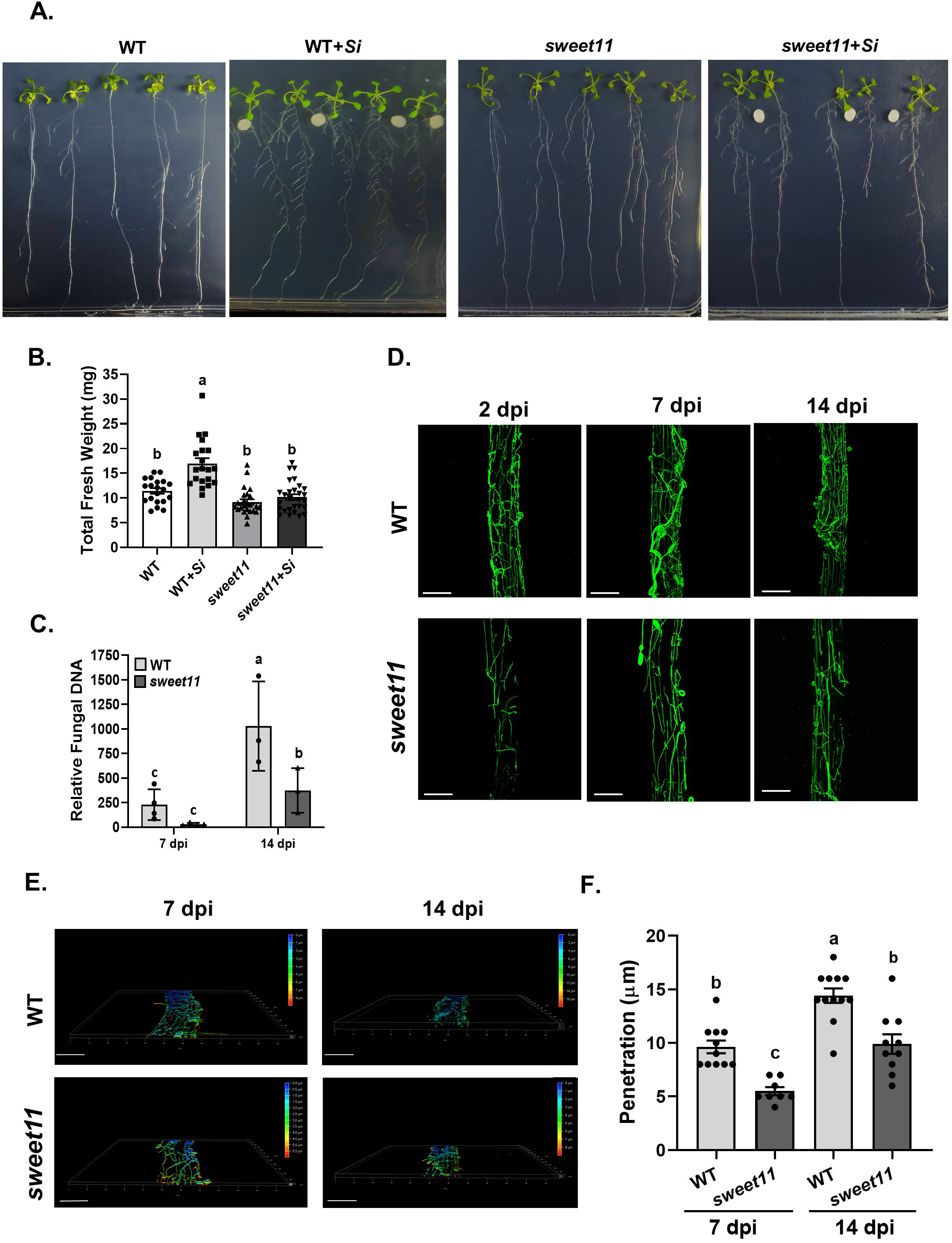
*S. indica* colonization with mutant *sweet11*: **A.** Representative picture of co-cultivation with WT, and *sweet11*. **B. Total fresh weight:** 7 days old Seedlings were co-cultivated with *S. indica* and total fresh weight was measured. Data represents mean ± SEM with n= >25. Different letters indicate significance difference after ANOVA analysis (*P* < 0.05, one-way ANOVA with Tukey’s test). **C. Relative Fungal DNA amount:** Roots at 14 dpi were collected and gDNA was isolated for RT-PCR using *SiTef1*. Colonization level in WT and *sweet11* mutant at 7 and 14 dpi. Asterisks indicate significant differences in *sweet11* mutant line (Different letters indicate significance difference (*P* < 0.05, Student’s *t*-test). Data represents mean ± SEM of independent biological replicates. Each sample contains roots from 10 seedlings. **D. Representative picture of Alexa Flour 488- WGA stained *S*. *indica* in WT and *sweet11* mutant roots:** WT and *sweet11* mutant roots were harvested and stained with fungus staining florescent dye Alexa Flour 488- WGA and pictures were taken under confocal microscope at 7 and 14 dpi. Scale bar = 100 μm. **E. Penetration Analysis of *S. indica* in plant roots:** Confocal images of *S. indica* colonization in wild-type (WT) and *sweet11* mutant roots were captured at 7 dpi (days post inoculation) and 14 dpi using Alexa Fluor 488-labeled wheat germ agglutinin (WGA) staining. The images were optically sectioned and stacked into a Z-plane, illustrating the depth of fungal penetration into the roots. Imaging began at the surface of the root (0 μm) to track the presence of fungal colonization. The red color represents the deepest zone of fungal penetration, where *S. indica* mycelia were detected last, while green and blue indicate shallower regions, with blue representing the root surface (Scale bar = 100 μm). **F. Statistical analysis of *S. indica* penetration in roots:** The graph shows comparative depth of root tissue colonization by *S. indica* in WT and *sweet11* at 7 and 14 dpi. Asterisks indicate significant differences in *sweet11* mutant (*P* < 0.05, one-way ANOVA with Tukey’s test). Data represents mean ± SD of at least 10 plant roots.

### 3.4. *sweet11* mutant show decreased *S. indica* colonization with less penetration in roots

To assess the impact of *SWEET11* loss-of-function on *S. indica* colonization in roots, we first quantified relative fungal DNA levels in *sweet11* mutant roots compared to WT roots. Our relative fungal DNA analysis revealed that *S. indica* colonization was reduced in *sweet11* mutant roots compared to WT at 14 dpi (**Fig. 2C**). Confocal microscopy revealed a reduction in Alexa Fluor 488–WGA-stained fungal structures, consistent with the trend observed in relative fungal DNA levels (**Fig. 2D**). Furthermore, analysis of fungal penetration depth using z-stack measurements from optically sectioned images of *S. indica*-colonized roots revealed reduced penetration in *sweet11* roots at 7 and 14 dpi compared to WT roots (**Fig. 2E-F**). These observations suggest that *SWEET11* is required for fungal establishment.

### 3.5. *sweet11* mutants have altered sugar dynamics during *S. indica* colonization

Soluble sugars translocated from shoots serve as a critical resource for *S. indica* multiplication within roots (Opitz et al. 2021). To investigate the impact of *SWEET11* loss-of-function on sugar dynamics during *S. indica* symbiosis, we quantified the levels of sugars in shoot and whole seedlings in WT and *sweet11* using GC-MS/MS analysis (**Fig. 3A-B; Fig. S6A-B)**. In whole seedlings, *S. indica* colonization (14 dpi) increased the levels of many sugars including sucrose, glucose, fructose, galactose, arabinose and maltose in WT plants (**Fig. S6A-B**). In *S. indica*-colonized whole *sweet11* plant, the contents of these sugars were found to be reduced as compared to *S. indica*-colonized WT and did not change compared to non-colonized *sweet11* plants (**Fig. S6A-B**). We separately quantified sugar content in shoots at 14 dpi and found reduced sugar content in shoot upon *S. indica* colonization in WT shoots (**Fig. 3A-B**). Surprisingly, this could not happen in the shoots of *sweet11* mutant upon *S. indica* interaction. For instance, fructose, galactose, *myo*-inositol, lactose, xylose and maltose were reduced upon *S. indica* colonization in WT shoots, but they remain unchanged in similar condition in the shoots of colonized *sweet11* compared to their non-colonized control shoots (**Fig. 3A-B**). In shoot, *S. indica* induces *SWEET11* which might be for increasing sugar flow towards roots (**Fig. 1**). These findings highlight the distinct roles of *SWEET11* in regulating sugar dynamics during *S. indica* colonization.

**Fig. 3.**
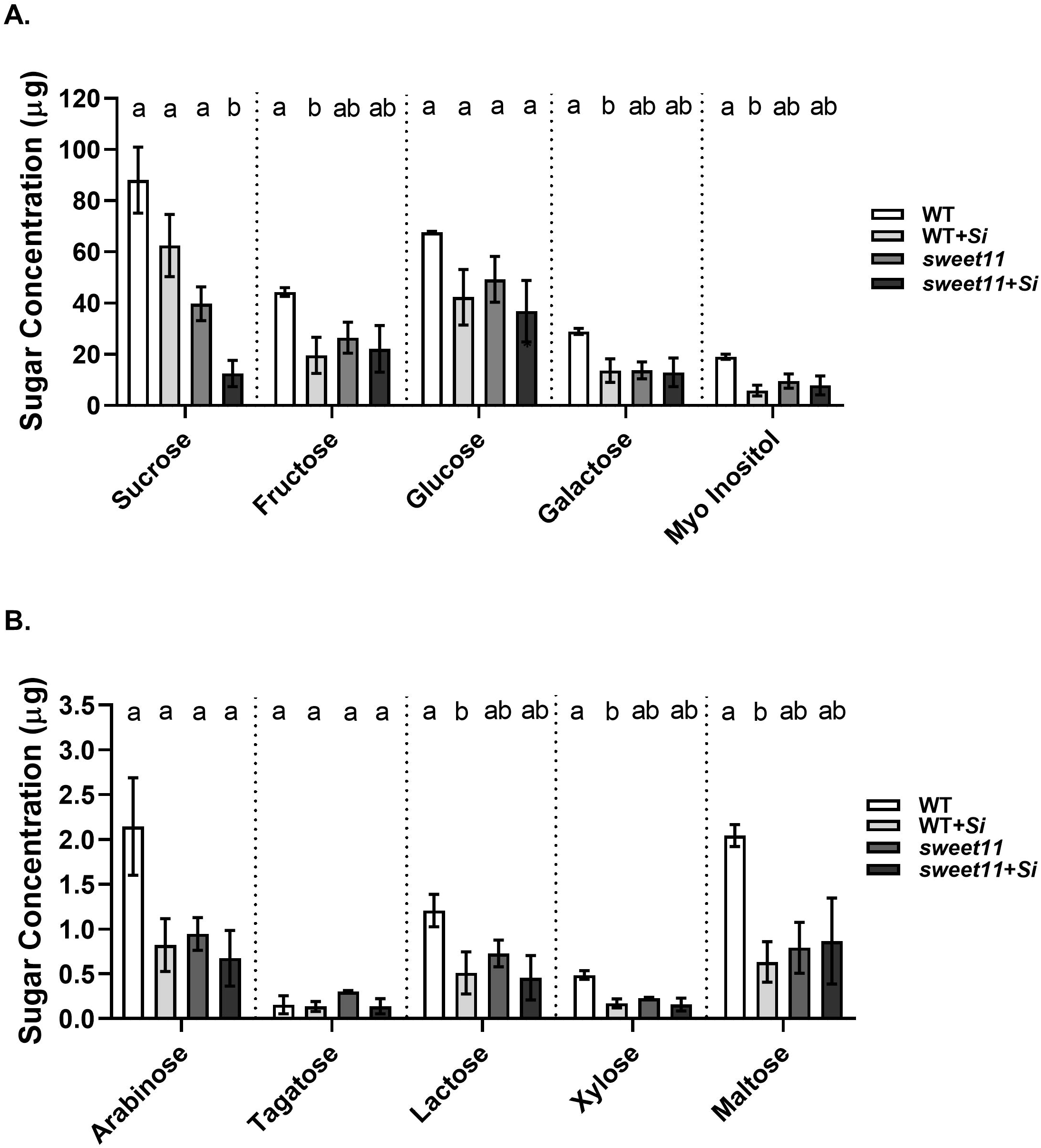
Sugar estimation in shoot by GC/MS analysis in WT and *sweet11*. **The content of major Sugar (A) and minor sugars (B) in WT and *sweet11* mutant shoots at 14 dpi:** 7 days old seedlings were co-cultivated and shoots were harvested and pooled at 14 dpi. Different sugars were estimated in lyophilized and powdered shoot material by GC/MS. Data represents mean ± SEM of 3 independent biological samples (3*50= 150). The different letters indicate significant differences in WT and *sweet11* mutant shoot upon *S. indica* colonization compared to their respective untreated control, as determined by ANOVA analysis (*P* < 0.05, one-way ANOVA with Tukey’s test).

### 3.6. Defense phytohormones are modulated in *sweet11* mutants

To observe the effect of SWEET11 loss on level of defense phytohormones we measured jasmonates (JA, JA-Ile), salicylic acid (SA) and abscisic acid (ABA) in non-colonized and *S. indica*-colonized seedlings at early (7 dpi) and late time points (30 dpi). In WT seedlings, *S. indica* colonization significantly induced the levels of both JA and JA-Ile, but no such increase was evident in *S. indica*-colonized *sweet11* mutant. (**Fig. 4A-B**). In *sweet11* mutants, basal JA and JA-Ile levels were found to be elevated compared to WT control (**Fig. 4A-B**). We observed the similar increase of SA levels in both WT and *sweet11* mutants upon *S. indica*-colonization at 7 dpi (**Fig. S7A**). At 30 dpi SA levels remain higher in *sweet11* mutants on *S. indica* colonization (**Fig. S7B**). However, ABA levels remain unchanged in WT upon *S. indica* colonization 7 and 30 dpi (**Fig. 4C, Fig. S7C**). However, in *sweet11* mutants, ABA levels were found to be significantly reduced in *S. indica*-colonized plants at 7 dpi (**Fig. 4C**). and 30 dpi (**Fig. S7C**). This suggests that SWEET11 loss-of-function might affect ABA level during *S. indica* interaction. These findings suggest that *SWEET11* might plays crucial role in modulating defense phytohormone levels, potentially influencing the level of *S. indica* colonization, plant immune responses and growth promotion.

**Fig. 4.**
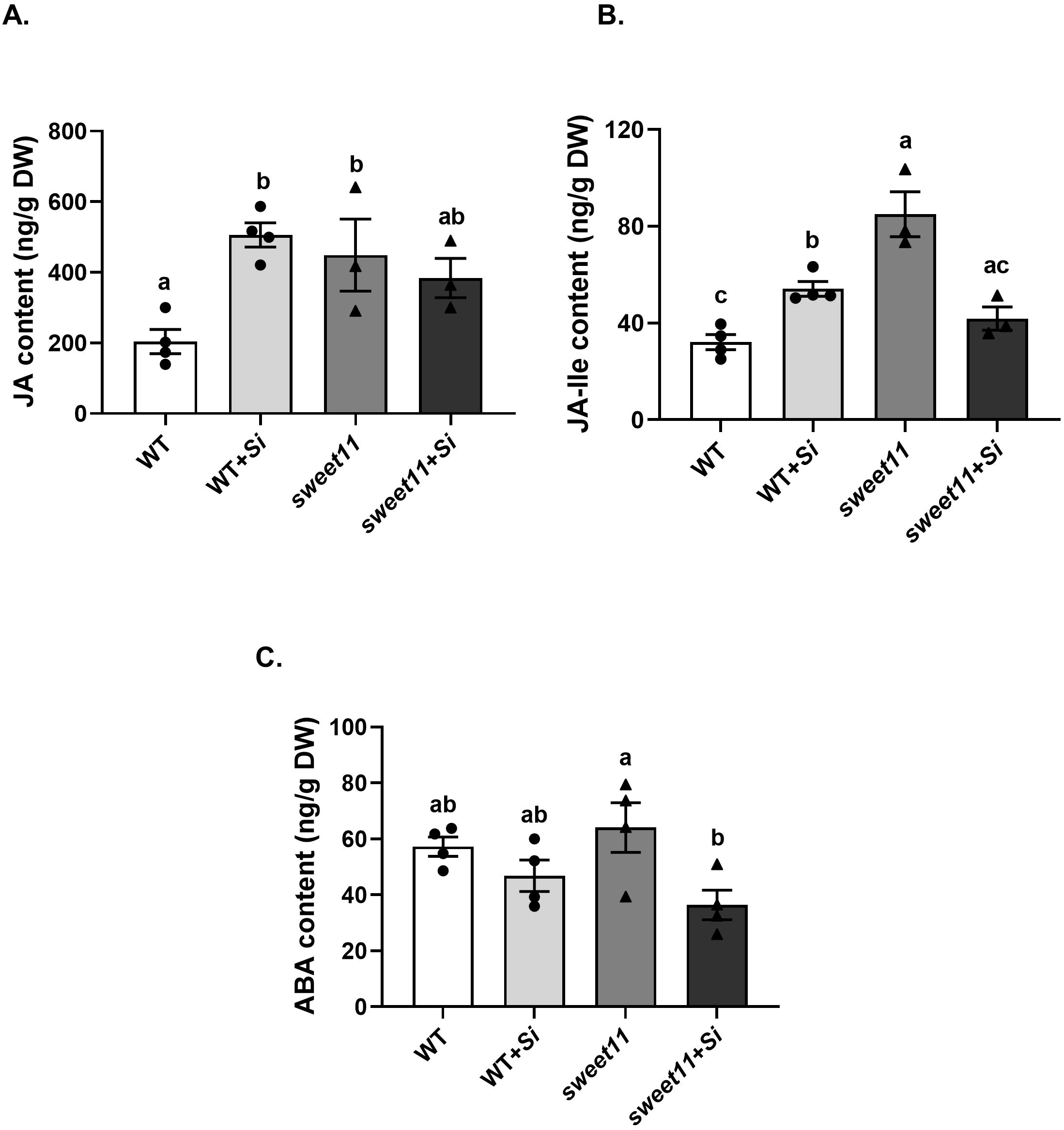
Defense phytohormones after *S. indica-*colonization. Levels of **A.** Jasmonic Acid (JA) **B.** Jasmonic acid-isoleucine (JA-Ile) and **C.** Abscisic acid (ABA) at 7 days of *S. indica* inoculation in WT and *sweet11*. The seedlings at 7 dpi were harvested, lyophilized, powdered and analysed by using LC-MS/MS. Bars are means ± SEM of at least three independent replicates (n=3*200=600). Different letters indicate significance difference after ANOVA analysis (*P* < 0.05, one-way ANOVA with Tukey’s test).

### 3.7. CT-induced SnRK2.8 interacts with the C-terminal of SWEET11

SWEET11 transporter interacts with SnRK2s which are ABA activated kinases (Chen et al. 2022). To investigate possible link of SWEET11 and SnRK2s in *S. indica* symbiosis, we explored the expression level of *SnRK2s* in response to *S. indica* colonization and cellotriose (CT) treatment in the previously published microarray and RNA sequencing data (Vahabi et al. 2015; Johnson et al. 2018). Among the *SnRK2s*, *SnRK2.8* stood out as one of the most highly induced genes during *S. indica* colonization. In our expression analysis, SnRK2.8 shows up to a 20-fold increase in expression at 14 dpi in roots (**Fig. 5A**). Interestingly, it shows up to 25-folds upregulation in *S. indica*-colonized plant’s shoot at 7 and up to 3.5-folds at 14 dpi (**Fig. 5B**). Moreover, our time-course analysis revealed that *SnRK2.8* transcripts peak at 4 hours post-CT treatment, with an 8-fold induction, before gradually declining in seedlings (**Fig. 5C**). The expression patterns of SWEET11 and SnRK2.8 showed a coordinated trend in both shoot and root tissues following *S. indica* and CT treatments, suggesting a potential functional association between these proteins. To examine whether this correlation reflects a direct physical interaction, we performed a series of interaction assays. Previous study has indicated that, under drought conditions, SnRK2 kinases can phosphorylate the C-terminal region of SWEETs, thereby enhancing sugar export and promoting root development (Chen et al. 2022). Based on this premise, we tested the interaction of SnRK2.8 with the C-terminal domains of SWEET11 and SWEET12 using a yeast two-hybrid (Y2H) system. Our Y2H analysis revealed a strong interaction between SnRK2.8 and the cytosolic C-terminal region of SWEET11 (SWEET11-CT). This interaction was clearly observed on both triple dropout (Leu/-Trp/-His; TDO). In contrast, no interaction was detected between SnRK2.8 and the C-terminal domain of SWEET12 (**Fig. 6A**). Importantly, none of the constructs exhibited autoactivation, confirming the specificity of the observed interaction. To validate these findings in a plant system, we conducted a split luciferase complementation imaging (LCI) assay. Consistent with the Y2H results, a strong luminescence signal confirmed the interaction between SnRK2.8 and SWEET11-CT *in planta* (**Fig. 6B-C**). Further confirmation was obtained through a biomolecular fluorescence complementation (BiFC) assay in *Nicotiana* leaves, where reconstituted fluorescence signals supported the physical association of SnRK2.8 with SWEET11-CT (**Fig. 6D**). For assay validation, appropriate controls were included: CML42–CNGC19 served as the negative control, while CAM2–CNGC19 was used as the positive control in both LCI and BiFC experiments (Meena et al. 2019). These controls performed as expected, reinforcing the reliability of our results. This interaction suggests that SnRK2.8 might regulate SWEET11 activity through phosphorylation. Reportedly, the expression of SWEET11 and SWEET12 were shown to be CORK1-dependent upon sucrose treatment (Gandhi et al. 2024). So, we also checked the interaction of the C-terminal domains of SWEET11 and SWEET12 with CORK1-kinase domain. Our observations suggested that there is no interaction of CORK1 with SWEET11 and SWEET12 (**Fig S8A-C**) which further emphasized SnRK2.8-mediated SWEET11 regulation in *S. indica*-symbiosis.

**Fig. 5.**
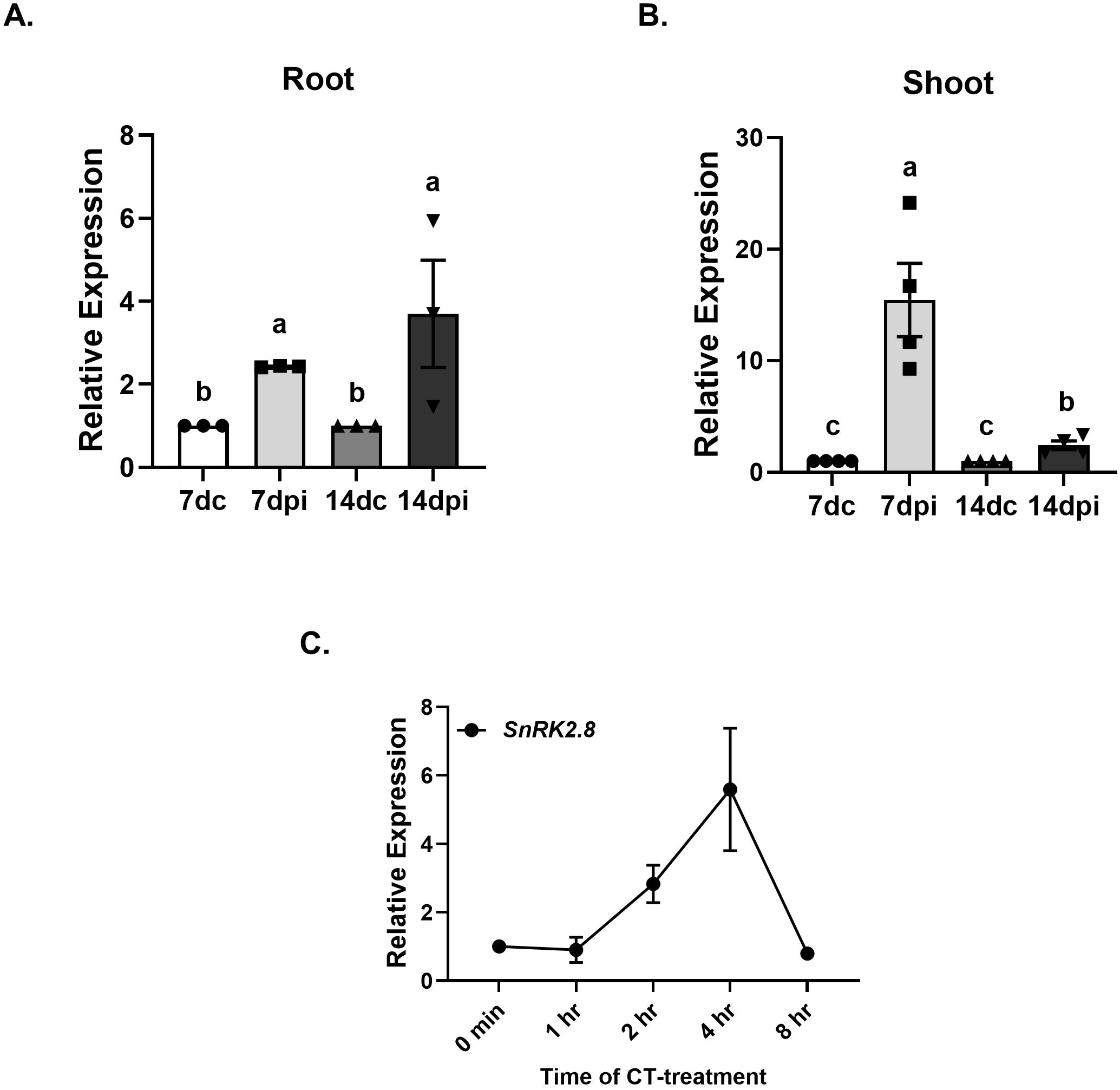
SnRK2.8 expression analysis. ***SnRK2.8* level in (A) shoot and (B) roots:** 7 days old seedlings were exposed to *S. indica* plugs and the relative expression of *SnRK2.8* was estimated post 7 and 14 dpi, separately in shoots and roots where non-treated control (7dc and 14dc) assumed as 1 fold. Data represents mean fold change ± SEM (N = 3-4, 3-4*10). **C. Time course expression analysis of *SnRK2.8* post CT-treatment:** 10 days old seedlings were exposed to 10 µM CT and *SnRK2.8* expression was checked at 0, 1 hr, 2 hr, 4 hr and 8 hrs post CT-treatment. Data represents mean fold change ± SEM (N = 4, 4*10). Different letters indicate statistically significant differences among samples as determined by using multiple *t*-tests (Holm-Sidak method), with *P* = 0.05.

**Fig. 6.**
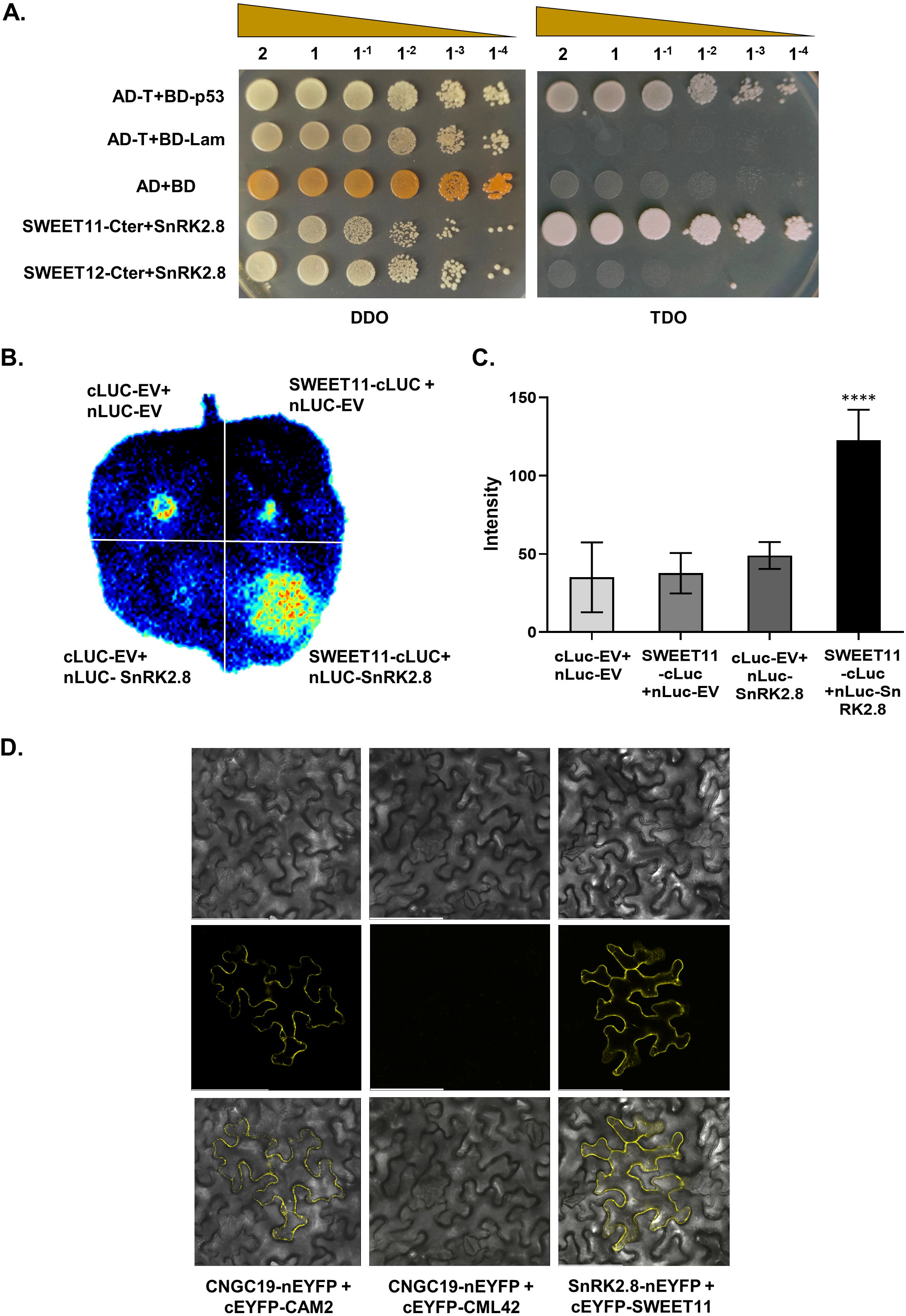
SnRK2.8 interaction analysis with SWEET11. **A. Yeast-Two-Hybrid assay:** SWEET11-Cter, SWEET12-Cter and full length SnRK2.8 were cloned in Y2H vectors and Y2H assay was performed. **B. Split luciferase assay:** This was performed using full length SnRK2.8 fused with nLUC and SWEET11 C-terminal (SWEET11-CT) fused with cLUC and cloned in the pCAMBIA1300 vector and transiently expressed in *N. benthamiana* by Agrobacterium-mediated infiltration. Images show only SWEET11-CT-cLUC+SnRK2.8-nLUC has stronger fluorescence due to positive interaction but not in cLUC-EV+cLUC-EV, SWEET11-cLUC+nLUC-EV and cLUC-EV+SnRK2.8-nLUC. **C. Quantification of mean intensity of signals:** SWEET11-CT-cLUC+ SnRK2.8-nLUC has significantly higher fluorescence compared to cLUC-EV+cLUC-EV, SWEET11-cLUC+nLUC-EV and cLUC-EV+SnRK2.8-nLUC. Data represents mean signal intensity ± SD (N = 10). Asterisks represent *P*-value **** >0.0001. D**. BiFC assay for SWEET11–SnRK2.8 interaction**. The assay was performed using SWEET11 fused with the C-terminal part of YFP and SnRK2.8 fused with the N-terminal part of YFP and transiently expressed in *Nicotiana benthamiana* by Agrobacterium-mediated infiltration. Images show YFP-mediated fluorescence derived from the protein–protein interaction. The brightfield image shows the plasma membrane. Scale bar = 100 µm. Interaction of CNGC19 with CML42 is shown as negative control and with CAM2 as a positive.

### 3.8. *snrk2.8* mutants exhibit impaired growth promotion during *S. indica* colonization

To further investigate the functional role of CT-induced and SWEET11-interacting SnRK2.8 in *S. indica* symbiosis, we analyzed growth promotion effect of *S. indica* in two *SnRK2.8* loss-of-function mutant lines i.e. *snrk2.8-1* and *snrk2.8-2*. After co-cultivation with *S. indica*, the growth assays revealed that both *snrk2.8* mutant lines displayed no growth promotion at 14 dpi compared to WT plants (**Fig. 7A-B**). Shoots and roots of *snrk2.8* mutants also do not show growth promotion compared to WT (**Fig. S9A-B**). Contrastingly, both *snrk2.8* mutant lines exhibited higher colonization levels than WT roots at 14 dpi (**Fig. 7C**). These findings indicate that SWEET11 interacting SnRK2.8 plays a critical role in *S. indica*-mediated growth promotion, similar to SWEET11 and acts in the same pathway.

**Fig. 7.**
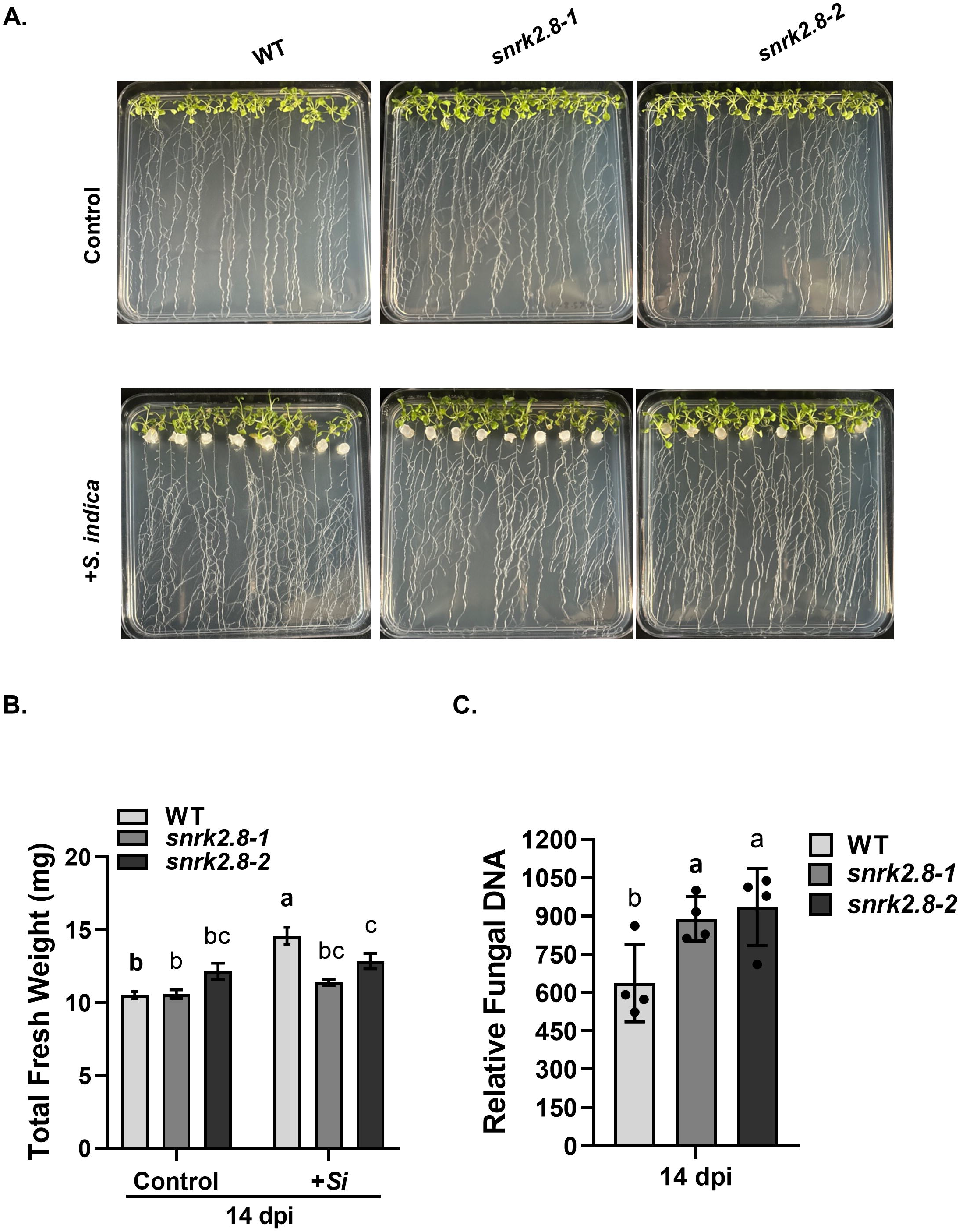
*S. indica* colonization with mutant *snrk2.8-1* and *snrk2.8-2*: **A.** Representative picture of co-cultivation with WT, *snrk2.8-1* and *snrk2.8-2* mutant captured at 14 dpi. **B. Total fresh weight:** 7 days old seedlings were co-cultivated with *S. indica* and total fresh weight was measured at 14 dpi. Data represents mean ± SEM with n ≥12. Different letters indicate significance difference after ANOVA analysis (**** *P* <0.0001, one-way ANOVA with Tukey’s test). **B. Relative Fungal DNA amount:** Roots at 14 dpi were collected and processed for genomic DNA isolation. RT-PCR was performed using *SiTef1* and *AtActin2* for estimating relative colonization level in WT and *snrk2.8* mutants at 14 dpi. Asterisks indicate significant differences in *snrk2.8* mutant lines over WT (Student’s *t*-test, *\* = P* < 0.05). Data represents mean ± SEM of four independent biological replicates (n = 4, 4*10).

### 3.9. RNA sequence data analysis indicates transcriptional reprogramming in *S. indica*-*sweet11* mutant roots

For gain more insight into the global transcriptional responses in *S. indica*-colonized roots (7 dpi) in the absence of SWEET11, we did RNA sequencing analysis. Transcriptome analysis revealed that the absence of SWEET11 causes extensive transcriptional reprogramming during *S. indica* colonization. At 7 dpi, WT roots exhibited 1,315 up- and 373 downregulated DEGs, whereas *sweet11* roots showed 564 up- and 212 downregulated DEGs (**Fig. 8A; Supplementary Data Sets 1–4**). DiVenn analysis demonstrated 955 DEGs unique to WT, 323 unique to *sweet11* with *S. indica*, and 1,466 unique to *sweet11* controls (**Fig. 8B**). A total of 313 DEGs were commonly upregulated in both WT and *sweet11* upon colonization, while 773 and 198 were uniquely induced in WT and s*weet11*, respectively (**Fig. 8C(a)**). Conversely, 182 and 161 DEGs were uniquely downregulated, with only 41 shared (**Fig. 8C (b)**). Notably, two common genes namely *RAP2.4D* (*RELATED TO AP2.4D*) and *SPIRAL1-LIKE2* (*SP1L2*) showed opposite regulation between genotypes, being upregulated in WT, but downregulated in *sweet11*. On contrary, three common genes namely *ISOCHORISMATE SYNTHASE 2* (*ICS2*), *PEROXISOMAL AND MITOCHONDRIAL DIVISION FACTOR 1* (*PMD1*) and *HYDROXYPYRUVATE REDUCTASE 1* (*HPR1*) were downregulated in WT, but upregulated in *sweet11* (**Fig. 8C(c)**). Differential expression of these genes indicates large scale transcriptional reprogramming in *S. indica*-colonized *sweet11* mutant compared to WT.

**Fig. 8.**
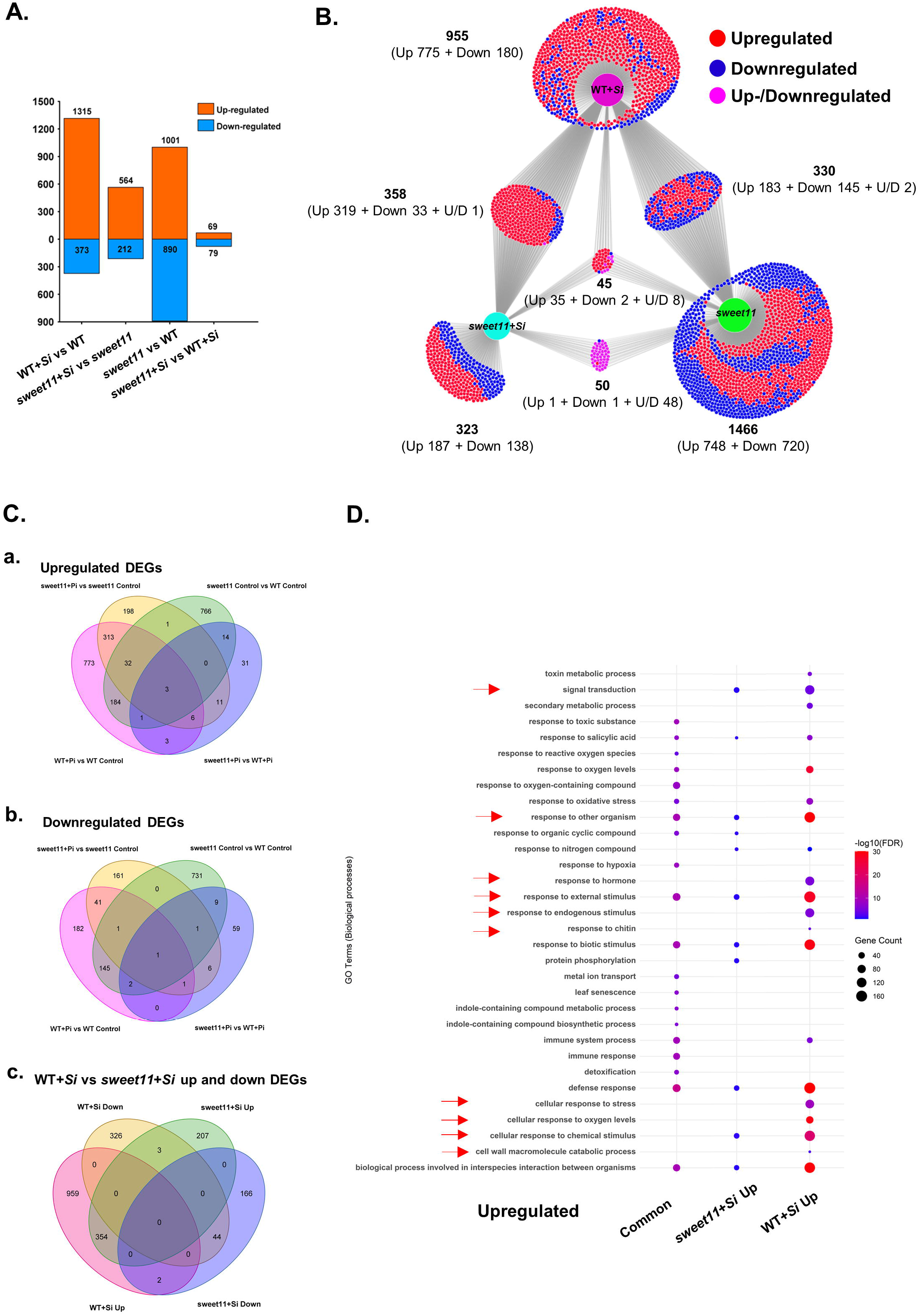
Transcriptomics analysis of *sweet11* control and *S. indica-*colonized *sweet11* mutant roots compared to WT. **A.** Numbers of differentially expressed genes (DEGs) in WT with *S. indica*, *sweet11* mutant and *sweet11* with *S. indica* at 7 dpi (Log2FC = ±0.58). **B.** Venn diagram showing the numbers of upregulated DEGs in WT with *S. indica*, *sweet11* mutant and *sweet11* with *S. indica* (Log2FC = +0.58). **C.** Venn diagram showing the numbers of downregulated DEGs in WT with *S. indica*, and *sweet11* with *S. indica* (Log2FC = -0.58). **D.** DEGs belonged to different biological processes upregulated in response to *S. indica* in roots at 7 dpi in WT and *sweet11* roots. The size and color of the circle in the graph represents number of genes and fold enrichment (-Log10FDR), respectively of corresponding biological process.

### 3.10. GO and clustering analyses reveals differential transcriptomic responses in *sweet11* mutant roots with and without *S. indica* colonization

Gene Ontology (GO) analysis of upregulated DEGs in *S. indica*-colonized WT and *sweet11* roots relative to their respective controls revealed that WT roots exhibited unique enrichment of biological processes, including toxin and secondary metabolism, responses to endogenous stimuli, hormones, and chitin, cellular responses to stress and oxygen, and cell wall macromolecule catabolic processes **(Fig. 8D)**. These processes are most likely associated with fungal perception, successful colonization, growth promotion, and enhanced sugar availability, and were less represented in *sweet11* under similar conditions. In GO analysis, *S. indica*-colonized *sweet11* roots exhibited reduced enrichment of upregulated DEGs in KEGG pathways and biological processes such as signal transduction, responses to other organisms, external and biotic stimuli, and defense responses compared to *S. indica*-colonized WT roots (**Fig. 8D, Fig. S10A-D**). Expression level analysis of the common up- and down-regulated DEGs of *S. indica*-colonized WT and *sweet11* roots has further revealed their expression less upregulated and more downregulated, respectively (**Fig. S11A and B**). Also, the expression of DEGs related to ABA signaling and biosynthesis such as *PYL*s (*PYL5, PYL6*), *SnRK2.7* and *SDR*s (*SDR1, SDR3*) shows reduction whereas DEGs related to SA (*ALD1*, *ICS2*, *NPR3*, *PAL2*, *PBS3* etc.), and JA (*JAZ1*, *LOX1*, and *LOX3*) show induction in *sweet11* compared to WT on colonization (**Fig. S12A-C**). Moreover, phytohormone defense-related WRKYs such as WRKY8, 18, 28, 30, 40, 46, and 70 were found to be comparatively less induced in *S. indica*-colonized *sweet11* than WT (**Fig. S12B**). Furthermore, DEGs related to perception, nutrient transport and calcium channels have also showed reduced expression patterns in *sweet11* with *S. indica* (**Fig. S13A-C**). Moreover, the DEGs comparison of *sweet11* over WT with *S. indica* colonization, pointed out important DEGs such as *PEN3*, *EDS1*, *MIOX4*, and *MRN1* which are significantly upregulated in *S. indica*-colonized *sweet11* roots and are mainly related to growth and defense (**Fig. 9C(a)**). All these observations suggest that SWEET11 loss-of-function has disrupted *S. indica*-induced nutrient transport, fungal perception, phytohormone signaling and accommodation of *S. indica* in plant roots.

**Fig. 9.**
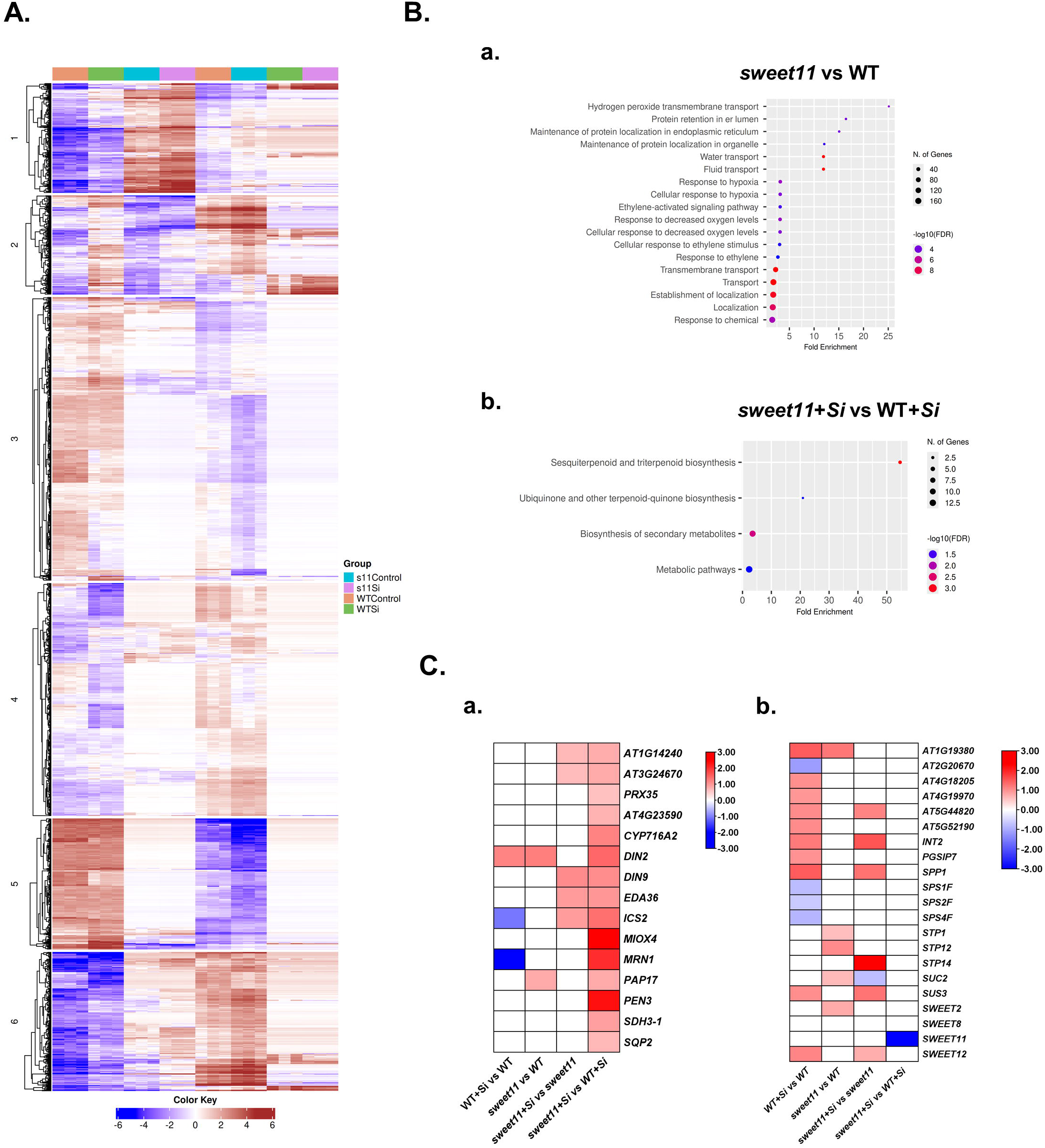
Clustering and KEGG pathway analysis of *S. indica-*colonized WT and *sweet11* mutant roots. A. Clustering: performed using raw read count data of all samples. All DEGs in *S. indica*-colonized *sweet11* roots and *S. indica*-colonized WT roots were as input in iDEP tool and k-Means clustering analysis was performed at -Log10FDR = 0.05. **B. Enrichment analysis: a.** KEGG enrichment analysis of upregulated DEGs in *S. indica*-colonized WT roots over its non-colonized control. **b.** KEGG enrichment analysis of upregulated DEGs in *sweet11* roots with *S. indica* over WT with *S. indica*. The size and color of the circle in the graph represents number of genes enriched and fold enrichment (-Log10FDR = 0.05), respectively of a particular corresponding KEGG pathway. **D. Heat map of DEGs: a.** Defense related DEGs in all conditions. **b.** Sugar transport and mobilization related DEGs in all conditions. Log2FC of DEGs were used to generate heat map.

For better insights, we performed cluster analysis of DEGs with similar expression patterns in all conditions. The clustering of root transcriptome data revealed that loss-of-function mutant *sweet11* shows DEGs enriched in 6 clusters. Cluster 1 showed enrichment of DEGs in KEGG pathways related to zeatin biosynthesis and plant-pathogen interaction, along with biological processes associated with responses to chemical and bacterial stimuli, stress, SA signaling, external biotic stimuli and other organisms, oxidative stress, reactive oxygen species (ROS), and organic cyclic compounds. In WT, this cluster shows suppression upon *S. indica* colonization but in *sweet11*, it was pre-induced and further induced upon *S. indica*-colonized roots (**Fig. 9A**). Cluster 2 was enriched for DEGs in KEGG pathways linked to phenylpropanoid and secondary metabolite biosynthesis, with associated biological processes including response to hypoxia, interspecies interactions, immune response, and immune system processes. This cluster shows induction in WT but not in *sweet11*. Importantly, Cluster 4 showed enrichment of biological processes related to disaccharide and oligosaccharide biosynthesis, photorespiration, and carbohydrate metabolism. This cluster show suppression in WT but not much changed in *sweet11*. Molecular function analysis further revealed enrichment of intracellular transport activities for iron, zinc, and nitrate/nitrogen in clusters 3 and 4 which shows significant changes in WT but not in *sweet11* (**Fig. 9A, Supplementary Data Set-5**). Furthermore, GO analysis of *sweet11* control over WT control shows enrichment of GOs such as fluid transport, response to hypoxia, response to ethylene and membrane transport (**Fig. 9B(a)**). Further, GO analysis of *S. indica* colonized *sweet11* shows enrichment of GOs such as metabolic pathways, biosynthesis of secondary metabolites, sesquiterpenoid, triterpenoid, ubiquinone and terpenoid-quinone which might affect phytohormone biosynthesis such as ABA and the level of defense molecules (**Fig. 9B(b)**). In *sweet11* with *S. indica*, sugar transport related DEGs were found to be repressed as compared to WT with *S. indica* (**Fig. 9C(b)**). The data shows that loss of growth promotion and reduced colonization in *sweet11* mutants as compared to WT is due to pre-induction of defense related pathways.

## 4. Discussion

Sugar transport and mobilization are frequent targets of microbial manipulation because carbohydrates represent the primary carbon source for both pathogenic and mutualistic microorganisms (Chu et al. 2006; Streubel et al. 2013). Pathogens commonly employ effector proteins to alter the expression of host sugar transporters, particularly members of the SWEET family, thereby redirecting host carbon toward infection sites (Chu et al. 2006; Breia et al. 2021). Similarly, in mutualistic interactions, microbial partners depend on host-derived sugars to sustain colonization and growth (Rani et al. 2021). The root-endosymbiont *Serendipita indica* has its own sugar transporters (*Si*HXTs) to uptake mainly monosaccharides from host (Rani et al. 2016; Raj et al. 2021). However, the identity of the plant transporters responsible for delivering sugars to the fungus and coordinating carbon allocation during mutualism has remained largely unknown. Recent studies have established that balanced sugar availability, mediated by the host, is essential for maintaining growth-defense homeostasis during *S. indica* symbiosis through coordinated regulation of SWEET transporters, invertases, and sucrose synthases (Opitz et al. 2021; Jogawat et al. 2026). In our previous study, we demonstrated that the root-specific transporter *At*SWEET12 functions as a sugar valve, restricting excessive carbon supply to the colonizing fungus and thereby maintaining a sustainable mutualistic interaction (Jogawat et al. 2026). The present study extends this model by identifying *At*SWEET11 as a systemically induced sugar transporter that regulates long-distance carbon allocation from shoots to roots during symbiosis.

Unlike SWEET12, which is induced locally in roots, *At*SWEET11 was specifically induced in shoots following root colonization by *S. indica*, and *sweet11* mutants failed to exhibit fungal-mediated growth promotion. Furthermore, *sweet11* plants showed significantly reduced fungal colonization and penetration, indicating that SWEET11 is required for efficient establishment of the mutualistic interaction. Transcriptome analysis further supports this conclusion, as *sweet11* displayed broad suppression of sugar transport-related genes together with induction of the galactose transporter STP14, suggesting compensatory alterations in carbohydrate metabolism. In addition, the *S. indica*-responsive marker gene AT1G58420 (Zecua-

Ramirez et al. 2025) failed to be induced in *sweet11* roots. Collectively, these observations suggest that SWEET11 functions upstream of a broader sugar mobilization program required for successful fungal colonization and host growth promotion. The systemic induction of *SWEET11* during *S. indica* symbiosis is consistent with observations in other plant species. In tomato, *S. indica* similarly induces *SlSWEET11b* expression in shoots upon root colonization (De Rocchis et al. 2022), suggesting that long-distance regulation of SWEET11 may be evolutionarily conserved. SWEET11 has also been implicated in diverse biotic interactions beyond symbiosis. Overexpression of *StSWEET11* in potato enhances *Phytophthora infestans* colonization through increased sucrose accumulation in the apoplast (Wu et al. 2024), whereas during leafminer (*Phthorimaea operculella*) infestation, *StSWEET11* is suppressed in damaged leaves but induced in undamaged systemic leaves to promote tolerance (Mao et al. 2025). These studies suggest that SWEET11 participates in dynamic source-sink adjustments that coordinate carbon allocation during both beneficial and antagonistic interactions rather than functioning solely as a susceptibility factor.

Accumulating evidence further indicates that SWEET11 integrates sugar transport with immune signaling. Hexose trimer, cellotriose (CT), a fungal cell wall-derived elicitor from *S. indica*, activates plant defense responses and induces *SWEET11* expression (Johnson et al. 2018; Gandhi et al. 2024). CT-induced reactive oxygen species (ROS) accumulation is significantly reduced in *sweet11* mutants, and induction of the defense-associated transcription factors WRKY40 and WRKY70 also depends on SWEET11 (Gandhi et al. 2024). Consistent with these observations, our transcriptome analysis revealed robust induction of several WRKY transcription factors, including WRKY30, WRKY33, WRKY51, WRKY62, and WRKY70, in wild-type roots following colonization, whereas only WRKY33, WRKY66, and WRKY70 remained induced in *sweet11*. These findings suggest that SWEET11 contributes not only to carbon allocation but also to transcriptional reprogramming associated with defense during symbiosis. Interestingly, SWEET11 appears to play a role distinct from SWEET12. Loss of SWEET12 or the cytosolic invertases CINV1 and CINV2 results in increased *S. indica* colonization owing to altered sugar pools and compromised defense responses (Opitz et al. 2021; Jogawat et al. 2026), whereas loss of SWEET11 reduces fungal colonization. Together, these contrasting phenotypes support a model in which SWEET12 locally regulates sugar availability within colonized roots, whereas SWEET11 systemically mobilizes carbon from shoots to sustain fungal establishment.

Our study further identifies SnRK2.8 as a potential upstream regulator of SWEET11 during symbiosis. The expression of *SnRK2.8* was induced by both CT treatment and *S. indica* colonization, and SnRK2.8 physically interacted with the C-terminal region of SWEET11, suggesting that SWEET11 activity may be regulated through phosphorylation. Consistent with this hypothesis, *snrk2.8* mutants exhibited compromised fungal-mediated growth promotion similar to *sweet11* mutants. Recent studies have shown that ABA-responsive SnRK2 kinases phosphorylate the C-terminal regions of SWEET transporters to enhance sugar transport toward roots during drought stress (Chen et al. 2022). This indicates that kinase-mediated activation of SWEET proteins may represent a conserved regulatory mechanism across different environmental conditions. Our findings therefore suggest that SnRK2.8 may activate SWEET11 to facilitate long-distance sugar transport from shoots to colonized roots during symbiosis.

Beyond regulating sugar transport, SnRK2.8 also plays important roles in systemic immunity. During systemic acquired resistance (SAR), SnRK2.8 promotes nuclear accumulation of NPR1 through phosphorylation and also phosphorylates the *Pseudomonas syringae* effector AvrPtoB, thereby stabilizing NPR1-mediated defense responses (Lee et al. 2015; Li et al., 2019; Lei et al. 2020; Saxena et al. 2025). NPR1 has likewise been shown to be essential for *S. indica*-induced systemic resistance, and *npr1* mutants display enhanced fungal colonization together with compromised SAR (Stein et al. 2008; Zhang et al. 2024). Interestingly, *snrk2.8* mutants also exhibit increased fungal colonization, suggesting that SnRK2.8 may integrate systemic immune signaling with carbon allocation during symbiosis. Furthermore, *sweet11* plants accumulated significantly lower ABA levels and exhibited reduced expression of ABA biosynthetic and signaling genes following *S. indica* colonization. ABA-dependent SnRK2 signaling regulates the balance between SnRK1 and TOR signaling, thereby controlling autophagy (Belda-Palazon et al. 2020). Because host autophagy is essential for successful *S. indica* colonization (Zecua-Ramirez et al. 2025) and *ZmSWEET11* has recently been linked to autophagy under salinity stress (Xing et al. 2026), SWEET11 may similarly coordinate sugar transport, ABA signaling, and autophagy during fungal symbiosis. Although this possibility remains speculative for the current study, it provides a framework for future investigation.

Collectively, our findings support a model in which root-derived fungal signals, including CT and potentially mobile fungal effectors, systemically induce *SnRK2.8* and *SWEET11* in shoots to promote long-distance sugar allocation toward colonized roots. In contrast to SWEET12, which functions locally as a sugar valve within roots, SWEET11 regulates source loading and long-distance carbon transport required to sustain fungal growth and host development. Loss of SWEET11 disrupts this coordinated source-sink relationship, alters sugar transporter expression, attenuates ABA-and WRKY-associated signaling, and ultimately compromises fungal colonization and plant growth promotion. Thus, SWEET11 and SWEET12 perform complementary but spatially distinct functions in optimizing carbon partitioning, immune regulation, and maintenance of the mutualistic interaction. We therefore propose that SWEET11 serves as a central hub linking systemic sugar transport with kinase-mediated signaling, phytohormone responses, and fungal colonization during *S. indica* symbiosis (**Fig. 10**).

**Fig. 10.**
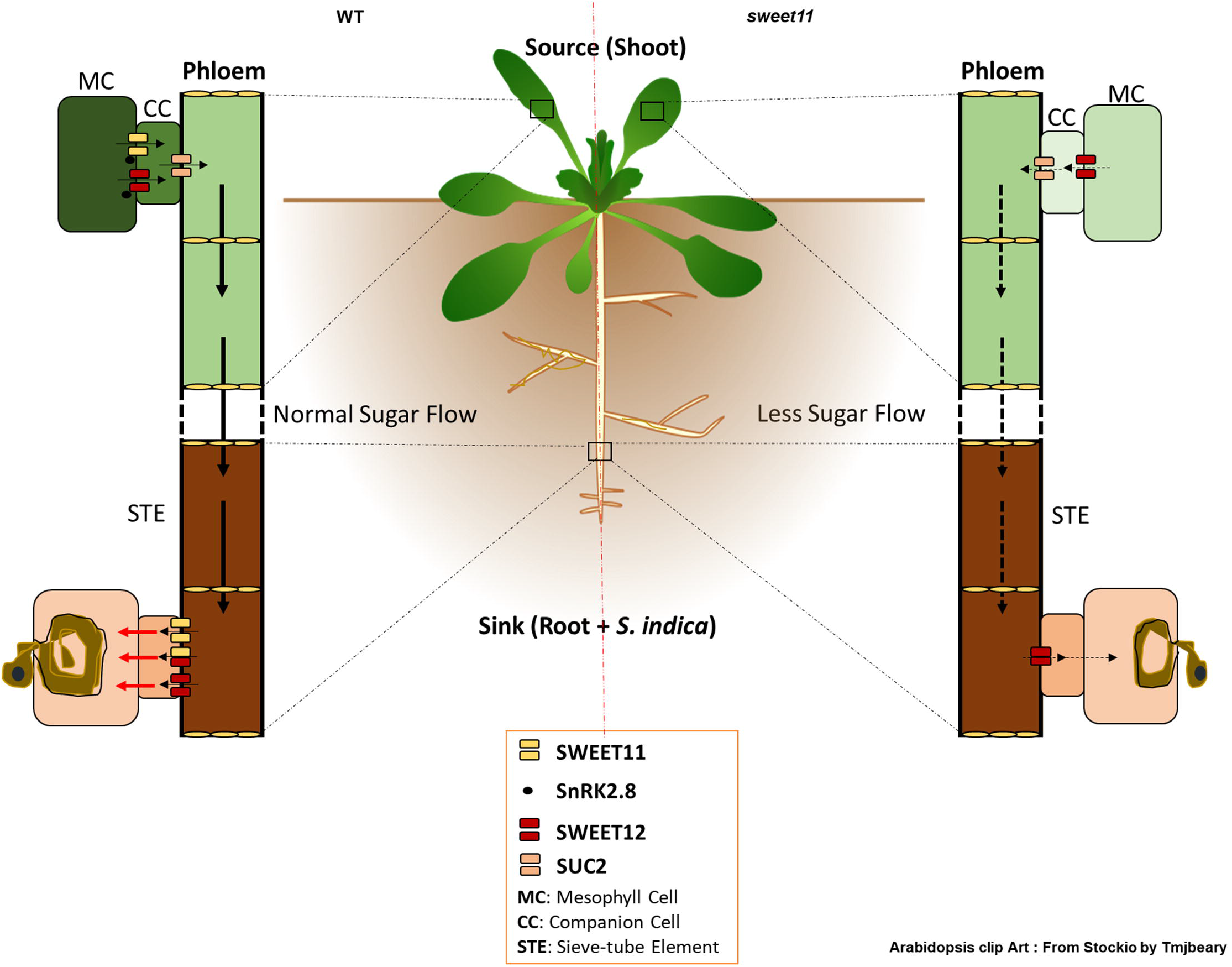
Proposed functional role of Arabidopsis SWEET11 during *S. indica* symbiosis. **Left Panel:** In WT plants upon *S. indica* colonization, sugar synthesis and mobilization is induced. In this symbiosis, SWEET11 functions like a “sugar releaser” for mobilizing sugars from shoot which supports colonization to a sufficient level for a successful symbiosis along with SWEET12. Due to this, SWEET11 functions as a crucial regulator of symbiosis between Arabidopsis and *S. indica*, maintaining fungal colonization level, plant immunity, and growth enhancement. **Right Panel:** In *sweet11* mutant, non-host resistance-like condition is observed, in which insufficient fungal level occurs to have no growth benefits to host and also high plant defense. The function of SWEET11 is regulated by SnRK2.8 for unloading more sugars towards colonized roots where SWEET11 might be regulated to provide carbon source to the colonized fungus. This event is missing in *sweet11* mutant which might cause reduced colonization and no growth promotion.

## Data Availability

The SRA metadata that support the findings of this study are available at NCBI under BioProject accession number PRJNA1381270 with SRA Data: SRR36479779, SRR36479778, SRR36479769, SRR36479768, SRR36479767, SRR36479766, SRR36479775, SRR36479774, SRR36479773, SRR36479772, SRR36479771, SRR36479770).

## Acknowledgements and funding

This work was supported through funding under MK Bhan-Young Researcher Fellowship (File No. HRD-16016/2/2023-AFS-DBT) of Department of Biotechnology, Government of India and NIPGR core grants from DBT. SHM and MS received Junior Research fellowship from MKBYRF. AMN acknowledges DST-INSPIRE. We acknowledge NIPGR for core grant, central instrumentation, metabolome facilities and DBT-eLibrary Consortium (DeLCON) for providing access to e-resources. We thank sincerely Dr. Senthil-Kumar Muthappa from BRIC-NIPGR for *sweet11* seeds, Dr. Cécile Vriet from University of Poitiers, France for p*SWEET11-*GUS seeds and Prof. Praveen K. Verma from Jawaharlal Nehru University, New Delhi for pCAMBIA1300-NLuc, pCAMBIA1300-CLuc, and modified-pENTR plasmids.

## Authors’ contributions

AJ and JV designed the experiments, analyzed the results and wrote the manuscript. AJ, SHM, MS, and AMN performed the experiments. DG performed GC-MS and LC-MS experiments. All authors approved the manuscript.

