## Supplementary information for "The systemically induced sugar transporter SWEET11 regulates growth-defense trade-offs during *Serendipita indica* symbiosis in Arabidopsis"

**Supplementary Table 1: List of primer pairs used in this study**

| S. N. | Primer name | Sequence | Experiment |
|---|---|---|---|
| 1 | SWEET1RTF | TGTGGTACTTCGTGGTTCGT | RT-PCR |
| 2 | SWEET1RTR | TTGCAGTGTCCCTAATGCAC | RT-PCR |
| 3 | SWEET2RTF | TGGTGTTTGCTGTGGTTGGA | RT-PCR |
| 4 | SWEET2RTR | CGGCGAGGCAAACATTGAAA | RT-PCR |
| 5 | SWEET3RTF | GTCGGCATCCTTCTCGAATCT | RT-PCR |
| 6 | SWEET3RTR | CTGTCGTTAAGCCGAACCCA | RT-PCR |
| 7 | SWEET4RTF | CATTGCCGGCATTTGTGGAA | RT-PCR |
| 8 | SWEET4RTR | TGCGCAATTTAGAACCGTGG | RT-PCR |
| 9 | SWEET5RTF | ATGGCGGTGGTGATCTTCTG | RT-PCR |
| 10 | SWEET5RTR | CATGACGGTGAGAGGAGCAG | RT-PCR |
| 11 | SWEET6RTF | ATGGTGCATGAACAGTTGAA | RT-PCR |
| 12 | SWEET6RTR | CAACCGATGTTGTTGACGACC | RT-PCR |
| 13 | SWEET7RTF | TTACGGACTACCAACGGTGC | RT-PCR |
| 14 | SWEET7RTR | TTTGGCGGCCACAATAAACG | RT-PCR |
| 15 | SWEET8RTF | CTTTGGTCTCTTTGCCGCAC | RT-PCR |
| 16 | SWEET8RTR | TGAACCACAGGGAGACCGTA | RT-PCR |
| 17 | SWEET9RTF | CCACGAAAACGGATTTGCCA | RT-PCR |
| 18 | SWEET9RTR | GCCTCACCGATCCTTCAACA | RT-PCR |
| 19 | SWEET10RTF | CTTTTTCGTCTGCCTTGCCC | RT-PCR |
| 20 | SWEET10RTR | ATCCATAGCATCGCGCTGAA | RT-PCR |
| 21 | SWEET11RTF | GGCACAGTTTCATCCCCTGA | RT-PCR |
| 22 | SWEET11RTR | GGAAGAGGACTGCTTGCCAT | RT-PCR |
| 23 | SWEET12RTF | TCTGTCTGCGTTTTTGCTGC | RT-PCR |
| 24 | SWEET12RTR | GAGCCATATGACCGCACTGA | RT-PCR |
| 25 | SWEET13RTF | CGTCAGTGTTTTCGCAGCTC | RT-PCR |
| 26 | SWEET13RTR | GTAGAAGAGCCACGTGACGG | RT-PCR |
| 27 | SWEET14RTF | ACTTCTACGTTGCGCTTCCA | RT-PCR |
| 28 | SWEET14RTR | ACCGTTTTGGGCTTCTCTGT | RT-PCR |
| 29 | SWEET15RTF | TCCTCCCCTCCAAGTCTCTG | RT-PCR |
| 30 | SWEET15RTR | AGCGTGAAGGGCATGTACTC | RT-PCR |

| | | | |
|---|---|---|---|
| 31 | SWEET16RTF | GCGATTGCGGGAACAAGAAC | RT-PCR |
| 32 | SWEET16RTR | CGTCACAACCGTTTTAATAGCTGA | RT-PCR |
| 33 | SWEET17RTF | ATTACGGCATCGTCACTCCC | RT-PCR |
| 34 | SWEET17RTR | TCGCTTCCACATCCACTGTT | RT-PCR |
| 35 | AtACTIN2RTF | TCAGATGCCCAGAAGTCTTGTTC | RT-PCR |
| 36 | AtACTIN2RTR | GTGGATTCCAGCAGCTTCCA | RT-PCR |
| 37 | SiTEF1RTF | TCGTCGCTGTCAACAAGATG | RT-PCR |
| 38 | SiTEF1RTR | GAGGGCTCGAGCATGTTGT | RT-PCR |
| 39 | SWEET11RGF | GGCACAGTTTCATCCCCTGA | RNA Genotyping |
| 40 | SWEET11RGR | GGAAGAGGACTGCTTGCCAT | RNA Genotyping |
| 41 | SnRK2.8RTF | TCATTGAAGAAGCTCGGAAA | RTPCR |
| 42 | SnRK2.8RTR | TCGAGATCCATACTGCTACT | RTPCR |
| 43 | SWT11 Cter_EcoR1F | GGAATTCGTGCTTGGTTTTGCTCTCGGTGC | Y2H Assay |
| 44 | SWT11 Cter_BamH1R | CGGGATCCTCATGTAGCTGCTGCGGAAGAGG | Y2H Assay |
| 45 | SWT12 Cter_EcoR1F | GGAATTCAAATACTGCAAAACGCCGTCGG | Y2H Assay |
| 46 | SWT12 Cter_BamH1R | CGGGATCCTCAAGTAGTTGCAGCACTGTTTCTAAC | Y2H Assay |
| 47 | SnRK2.8EcoRIF | CCGGAATTCATGGAGAGGTACGAAATAGTGAA | Y2H Assay |
| 48 | SnRK2.8RBamHIR | CGCGGATCCTCACAAAGGGGAAAGGAGATCA | Y2H Assay |
| 49 | CORK1_Y2H_EcoR1_F | GGAATTCCTGAACGACGGAAGAGAGAT | Y2H Assay |
| 50 | CORK1_Y2H_BamH1_R | CGGGATCCTCAATGTCGTCGTCCATGTT | Y2H Assay |
| 51 | SWT11CterBamHIF | CGCGGATCCATGTACAAATACTGTAAAACGTCGC | Split Luciferase Assay |
| 52 | SWT11CterSalIR | ACGCGTCGACTCATGTAGCTGCTGCGGAA | Split Luciferase Assay |
| 53 | S12LUCCterBamHIF | CGCGGATCCATGTACAAATACTGCAAAACGCC | Split Luciferase Assay |
| 54 | S12LUCCterSalIR | ACGCGTCGACTCAAGTAGTTGCAGCACTGTT | Split Luciferase Assay |
| 55 | SnRK2.8nLUCBamHIF | CGCGGATCCATGGAGAGGTACGAAATAGTGAAGG | Split Luciferase Assay |

| | | | |
|---|---|---|---|
| 56 | SnRK2.8nLUCSalIR | ACGCGTCGACTCACAAAGGGGAAAGGAGATCAGC | Split Luciferase Assay |
| 57 | S11pENTRnsNotIF | ATAAGAATGCGGCCGCATGAGTCTCTTCAACACTGA | Cloning in pENTR vector |
| 58 | S11pENTRnsAscIR | TTGGCGCGCCTGTAGCTGCTGCGGAA | Cloning in pENTR vector |
| 61 | SnRK2.8NotIF | AAGGAAAAAAGCGGCCGCATGGAGAGGTACGAAATAGTGAAGG | Cloning in pENTR vector |
| 62 | SnRK2.8AscIR | TTGGCGCGCCTCACAAAGGGGAAAGGAGATCAGC | Cloning in pENTR vector |
| 63 | S11CBiFCAttBF | GGGGACAAGTTTGTACAAAAAAGCAGGCTTCATGTACAAATACTGTAAAACGTCGC | Gateway cloning in pSITE vector |
| 64 | S11CBiFCAttBR | GGGGACCACTTTGTACAAGAAAGCTGGGTTCTGTAGCTGCTGCGGAAGAG | Gateway cloning in pSITE vector |
| 65 | SnRK2.8BiFCAttBF | GGGGACAAGTTTGTACAAAAAAGCAGGCTTCATGGAGAGGTACGAAATAGTGAAGG | Gateway cloning in pSITE vector |
| 66 | SnRK2.8BiFCAttBR | GGGGACCACTTTGTACAAGAAAGCTGGGTTCCAAAGGGGAAAGGAGATCAGC | Gateway cloning in pSITE vector |
| 67 | TDNALBF | ATTTTGCCGATTTCGGAAC | T-DNA confirmation |

### Supplementary Figures:

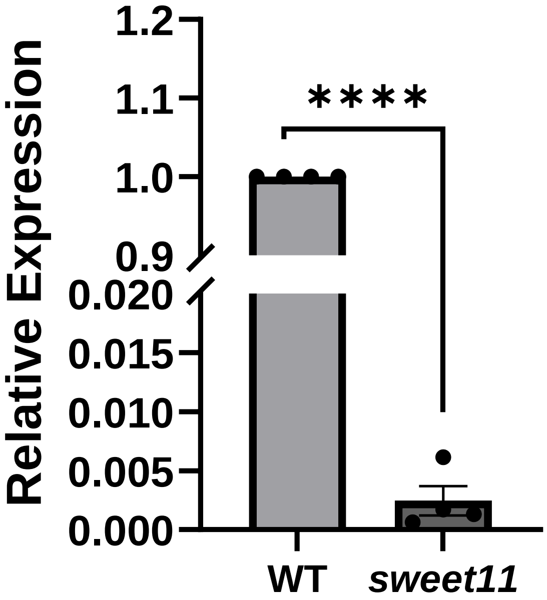


**Fig. S1. RNA genotyping of *sweet11* mutant line (SALK_073269):** From ten days old seedlings, RNA was extracted, cDNA was prepared and *SWEET11* transcripts level was checked by using *SWEET11* real time-PCR primers. Data represents mean fold change ± SEM of n = 4 (4*10=40). Asterisks indicate significant difference after Student's *t*-test, $P < 0.0001$.

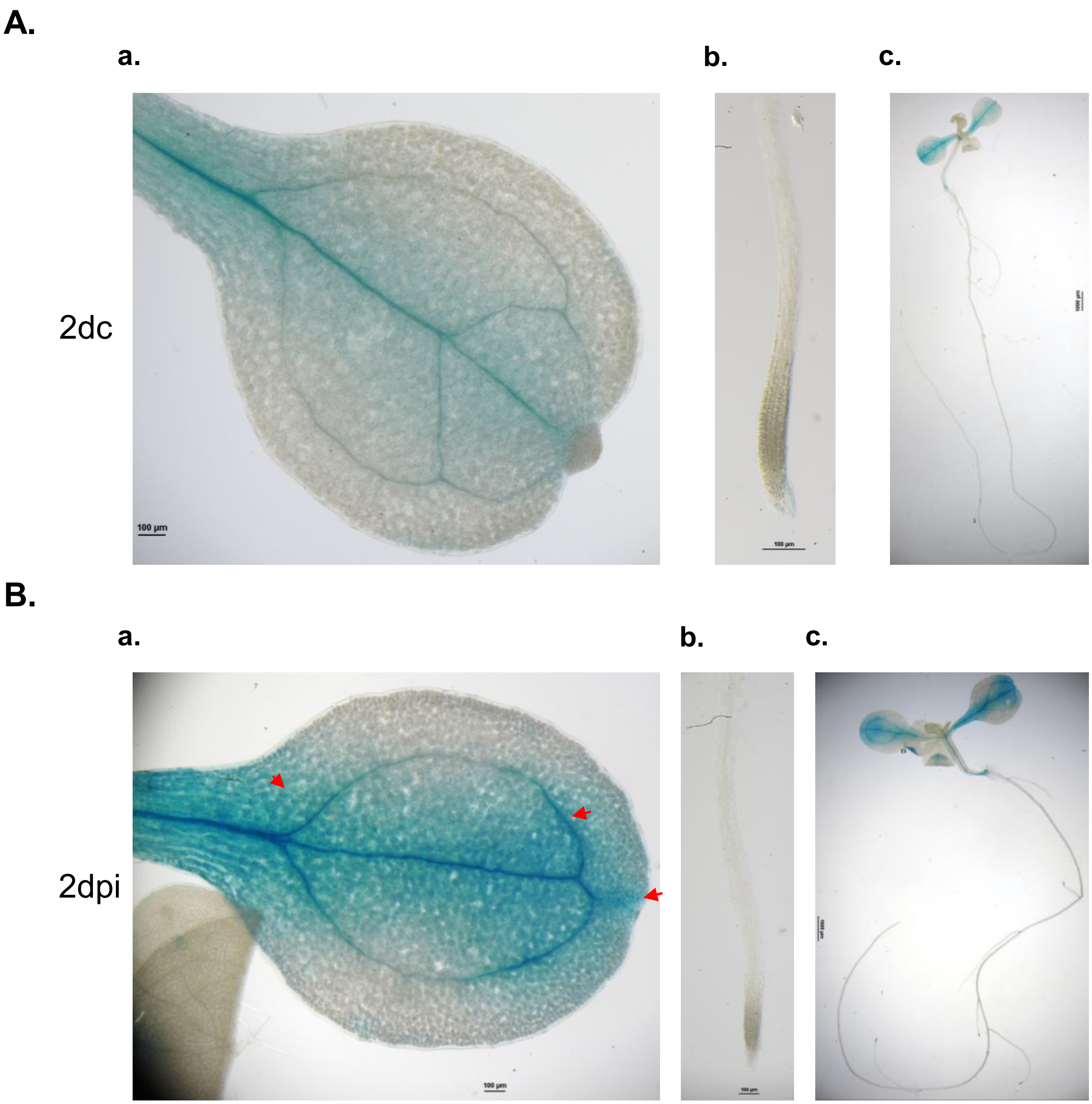


**Fig. S2. *SWEET11* promoter activity during *S. indica* symbiosis at 2 dpi.** p*SWEET11-GUS* expressing seedling were grown for 10 days on MS and then co-cultivated with *S. indica* on 1X PNM media. At 2 dpi, GUS assay was performed to check the activity of p*SWEET11-GUS*. The blue color shows *SWEET11* promoter activity at 2 dpi in the shoot and root. Images of leaf (**a**), root tip (**b**) and whole seedling (**c**) were captured in absence at 2dc (**A**) and presence at 2 dpi (**B**) of *S. indica*. Arrows indicate the induced GUS expression in different regions of seedling upon *S. indica* colonization compared to non-inoculated seedlings. Scale bar = 100 µm.

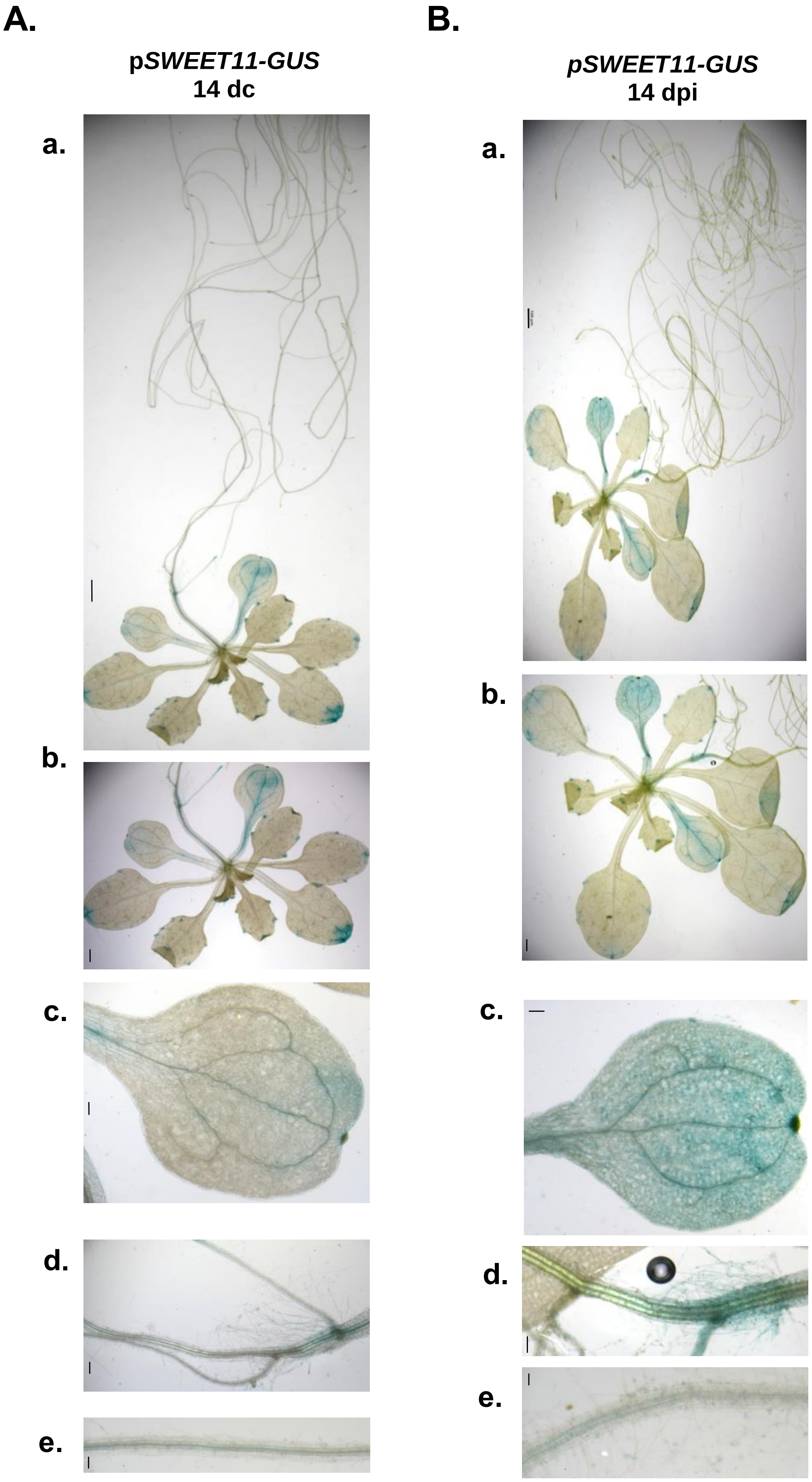
A.
pSWEET11-GUS
14 dc
a.
b.
c.
d.
e.
B.
pSWEET11-GUS
14 dpi
a.
b.
c.
d.
e.

**Fig. S3. *SWEET11* promoter activity during *S. indica* symbiosis at 14 dpi.** p*SWEET11-GUS* expressing seedling were grown for 10 days on MS and then co-cultivated with *S. indica* on 1X PNM media. At 14 dpi, GUS assay was performed to check the activity of p*SWEET11-GUS*. The blue color shows *SWEET11* promoter activity at 14 dpi in the shoot and root. Images of whole seedling (a), rosette (b), single leaf (c), root-shoot junction (d) and root (**e**) were captured in absence at 14dc (**A**) and presence at 14dpi (**B**) of *S. indica*. Arrows indicate the induced GUS expression in different regions of seedling upon *S. indica* colonization compared to non-inoculated seedlings (Scale bar = 100 μm).

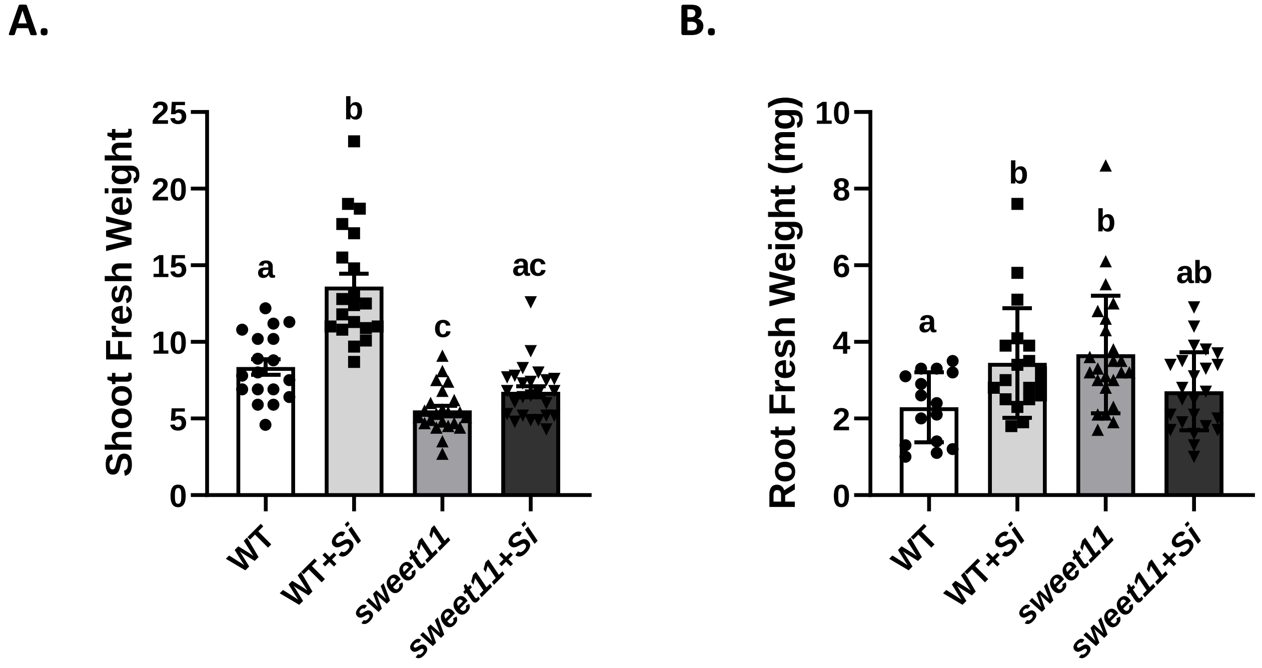


**Fig. S4. Growth assay of WT and *sweet11* after *S. indica* co-cultivation. A. Shoot fresh weight:** Seven days old seedlings were co-cultivated with S. *indica* and total fresh weight was measured at 14 dpi. Data represents mean ± SEM (n ≥ 18). **B. Root fresh weight:** Data represents mean ± SEM (n ≥ 15). Different letters indicate significance difference after ANOVA analysis ($P \leq 0.05$, one-way ANOVA with Tukey's test).

A.

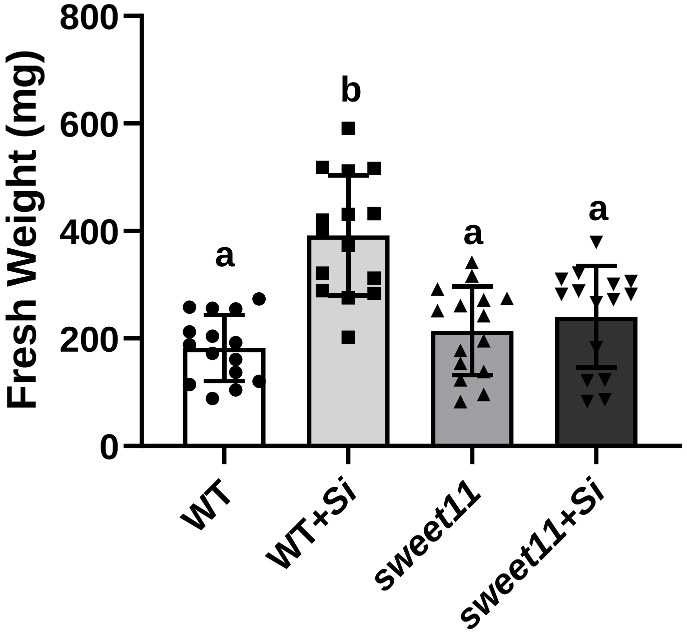


B.

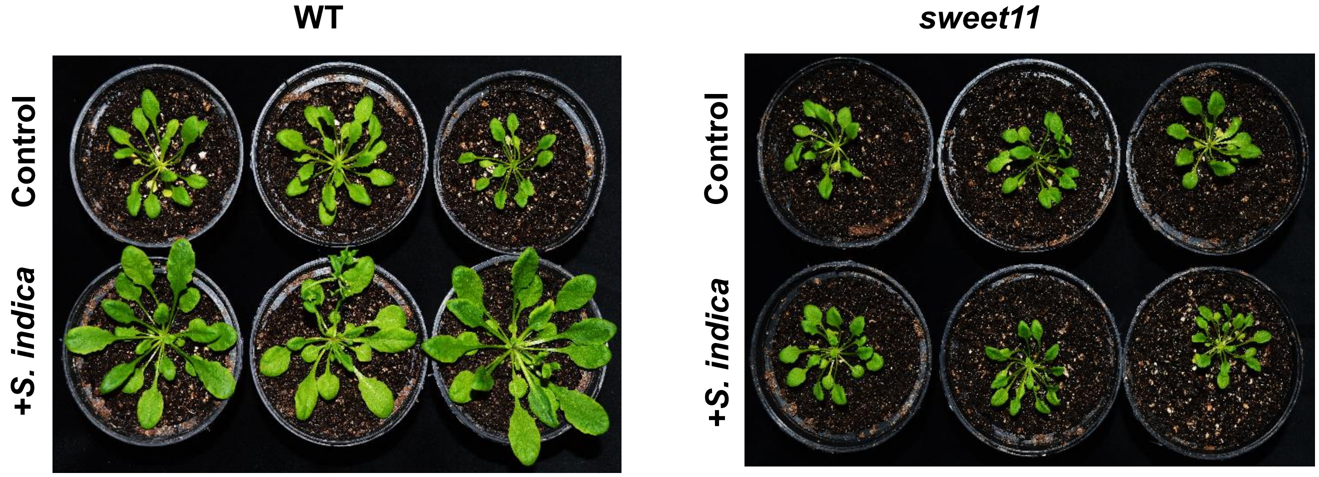


**Fig. S5. *S. indica* co-cultivation with WT and *sweet11* in soil. A. Fresh weight:** Two weeks old days old Seedlings were co-cultivated with *S. indica* and rosette fresh weight was measured at 30 dpi. Data represents mean ± SEM with n= 15. Different letters indicate significance difference after ANOVA analysis ($P \leq 0.05$, two-way ANOVA with Tukey's test). **B.** Representative picture of soil co-cultivation of *S. indica* with WT and *sweet11* mutant.

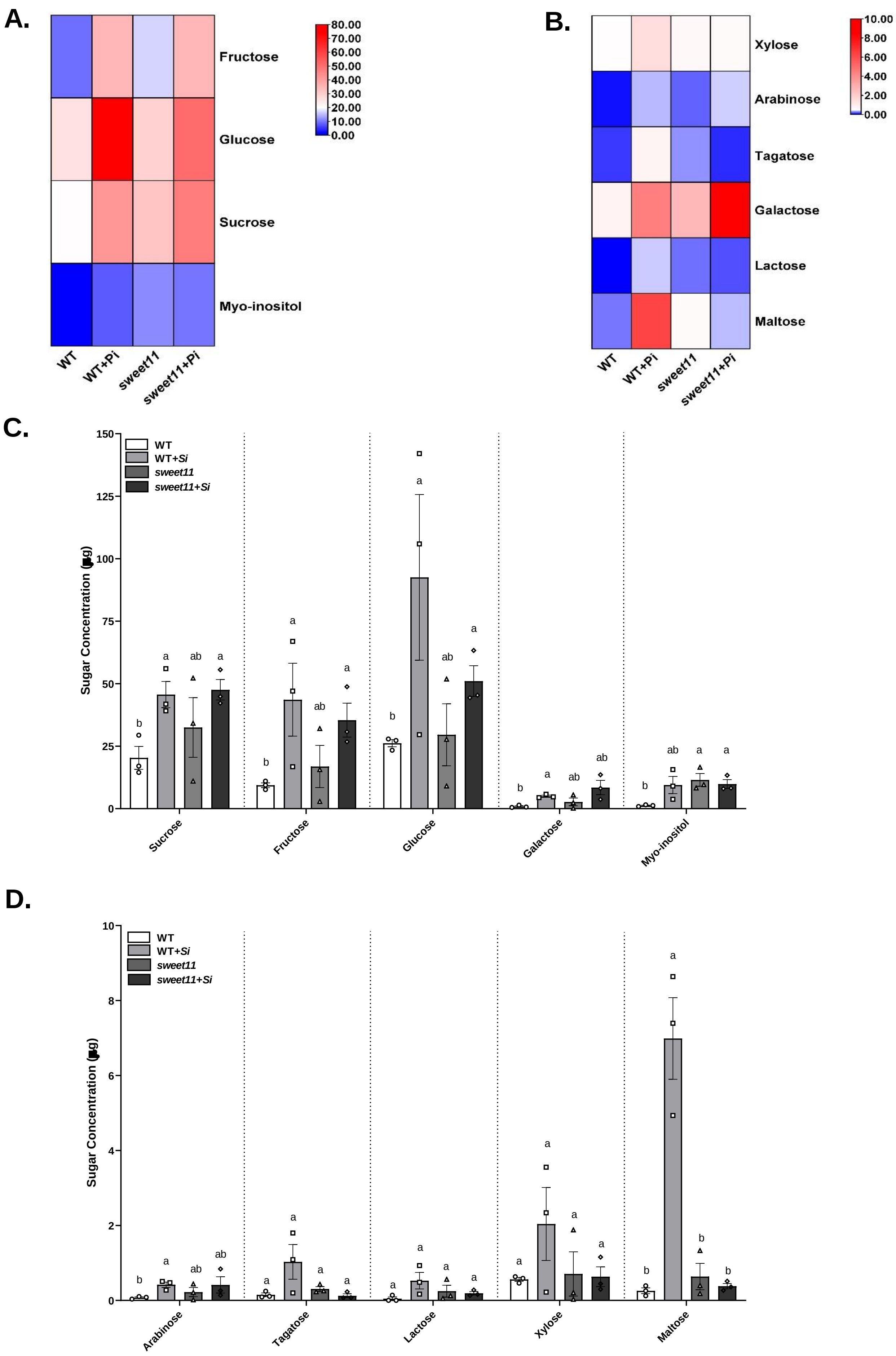
A.
Fructose
Glucose
Sucrose
Myo-inositol
WT
WT+Pi
sweet11
sweet11+Pi
80.00
70.00
60.00
50.00
40.00
30.00
20.00
10.00
0.00
B.
Xylose
Arabinose
Tagatose
Galactose
Lactose
Maltose
WT
WT+Pi
sweet11
sweet11+Pi
10.00
8.00
6.00
4.00
2.00
0.00
C.
WT
WT+Si
sweet11
sweet11+Si
Sugar Concentration
150
125
100
75
50
25
0
Sucrose
Fructose
Glucose
Galactose
Myo-inositol
D.
WT
WT+Si
sweet11
sweet11+Si
Sugar Concentration
10
8
6
4
2
0
Arabinose
Tagatose
Lactose
Xylose
Maltose

**Fig. S6. Sugar estimation in whole plant by GC/MS analysis. A, B. Sugar content in WT and *sweet11* mutant whole plants at 14 dpi:** Different sugars were estimated in lyophilized and powdered plant material harvested at 14 dpi when symbiosis is well established. Heat map data represents mean of 3 independent biological samples (3*50= 150). **C, D. Sugar content in WT and *sweet11* mutant whole plants at 14 dpi:** Data are presented as mean ± SEM. Different letters indicate statistically significant differences among samples as determined by using multiple *t*-tests (Holm-Sidak method), with $P = 0.05$.

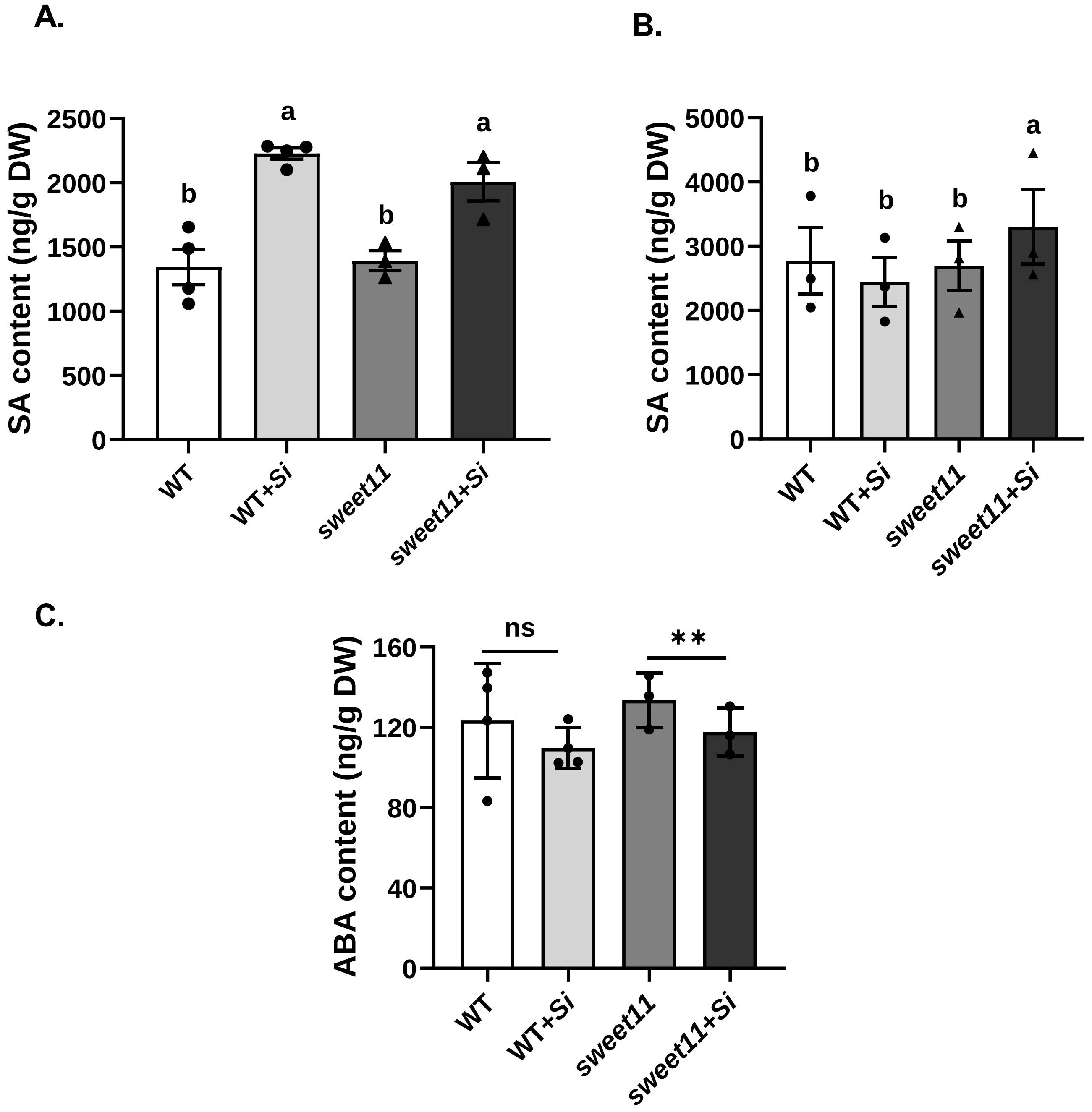

**Fig. S7. Salicylic acid and Abscisic acid content in *S. indica*-colonized seedlings. A. SA content at 7 dpi:** Seven days old seedlings of WT and *sweet11* seedlings were co-cultivated with *S. indica.* The samples were harvested at 7 dpi and lyophilized for LC-MS/MS analysis. Data are means ± SE of three independent replicates (n= 4*200= 800). **B. SA** and **C. ABA content at 30 dpi:** *S. indica* was co-cultivated with two week old seedlings of WT and *sweet11* seedlings*.* The rosette were harvested at 30 dpi and lyophilized for LC-MS/MS analysis. Data are means ± SE of 3-4 independent replicates. Each replicate has four rosettes. Different letter indicates significant difference calculated by one-way ANOVA analysis with Tukey's test with $P \leq 0.05$. Asterisks indicate significant difference after Student's *t*-test, $P < 0.001$.

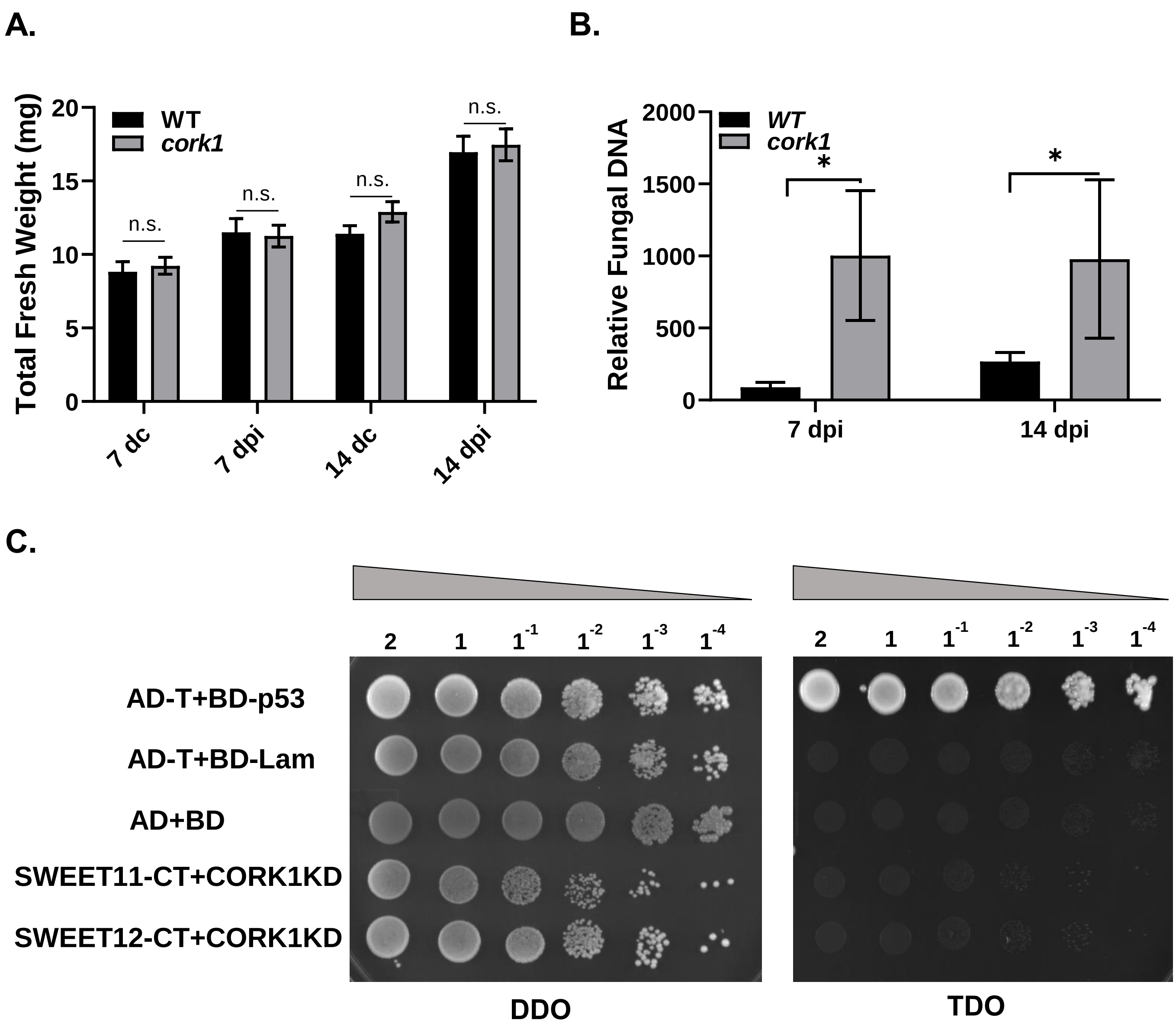

**Fig. S8. *cork1* mutant *S. indica* colonization and CORK1-KD interaction study with SWEETs. A. Total fresh weight:** 10 days old seedlings were co-cultivated with *S. indica* and total fresh weight was measured at 7 and 14 dpi (n ≥ 20). The n.s. indicate no significant differences in the treatment over its corresponding control (Student's *t*-test, n ≥ 20). **B. Relative Fungal DNA amount:** Roots at 7 and 14 dpi were collected and genomic DNA were isolated for RT-PCR using *SiTef1* as fungal marker and *AtActin2* as plant marker. Colonization level in Col-0 and *cork1* roots at 7 and 14 dpi in co-cultivation media was determined. Asterisks indicate significant differences in the treatment over its corresponding control (* $P < 0.05$, Student's *t*-test, n ≥ 3). Data represents mean ± SEM of at least 3 independent biological replicates. Each sample contains roots from 10 seedlings. **C. Yeast-Two-Hybrid assay:** SWEET11-Cter, SWEET12-Cter and CORK1-Kinase Domain (CORK1KD) were cloned in Y2H vectors and Y2H assay was performed.

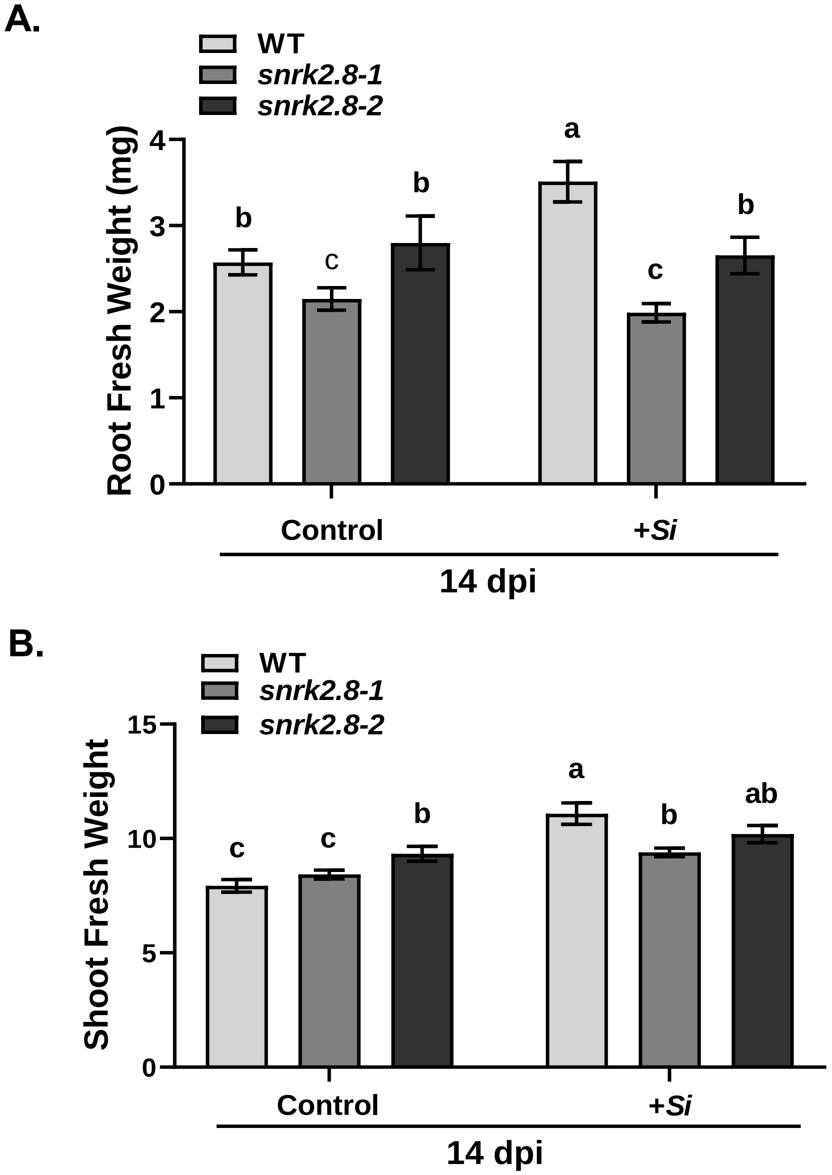


**Fig. S9. Root and shoot fresh weight of *snrk2.8* mutants:** 7 days old seedlings were co-cultivated with *S. indica* and fresh weights of root (**A**) and shoot (**B**) were measured separately. Data represents mean ± SEM with n ≥12. Different letters indicate significance difference after ANOVA analysis ($P$ <0.05, one-way ANOVA with Tukey's test).

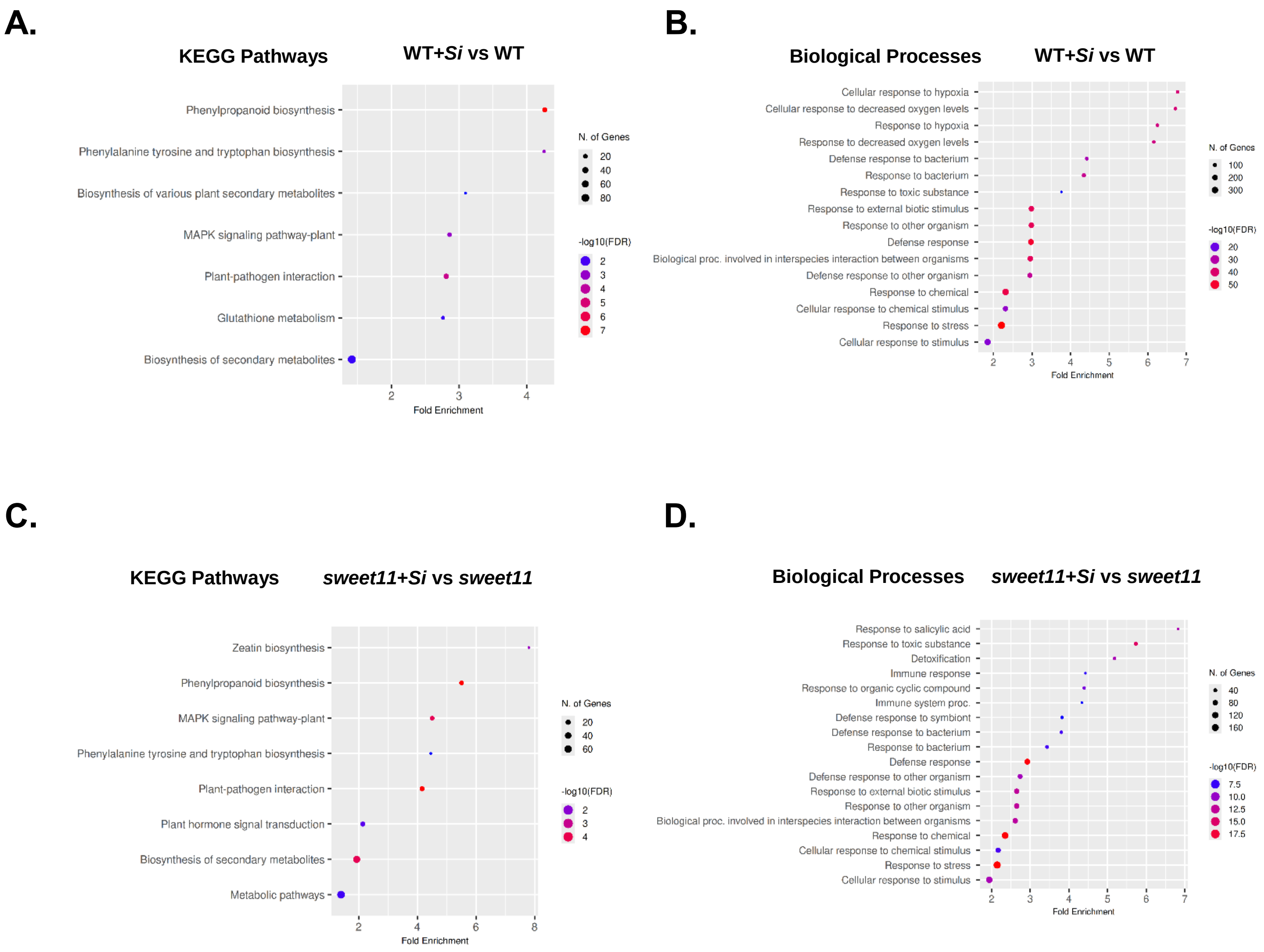


**Fig. S10. Gene ontologies enrichment analysis: A. KEGG pathway analysis of *S. indica*-colonized WT and *sweet11* mutant roots. A.** KEGG enrichment analysis of upregulated DEGs in *S. indica*-colonized WT roots over its non-colonized control. **B.** Biological processes enrichment analysis of upregulated DEGs in *S. indica*-colonized WT roots over its non-colonized control. **C.** KEGG enrichment analysis of upregulated DEGs in *S. indica*-colonized *sweet11* roots over its non-colonized control. **D.** Biological processes enrichment analysis of upregulated DEGs in *S. indica*-colonized *sweet11* roots over its non-colonized control. The size and color of the circle in the graph represents number of genes enriched and fold enrichment (-Log10FDR = 0.05), respectively of a particular corresponding KEGG pathway or bioprocess.

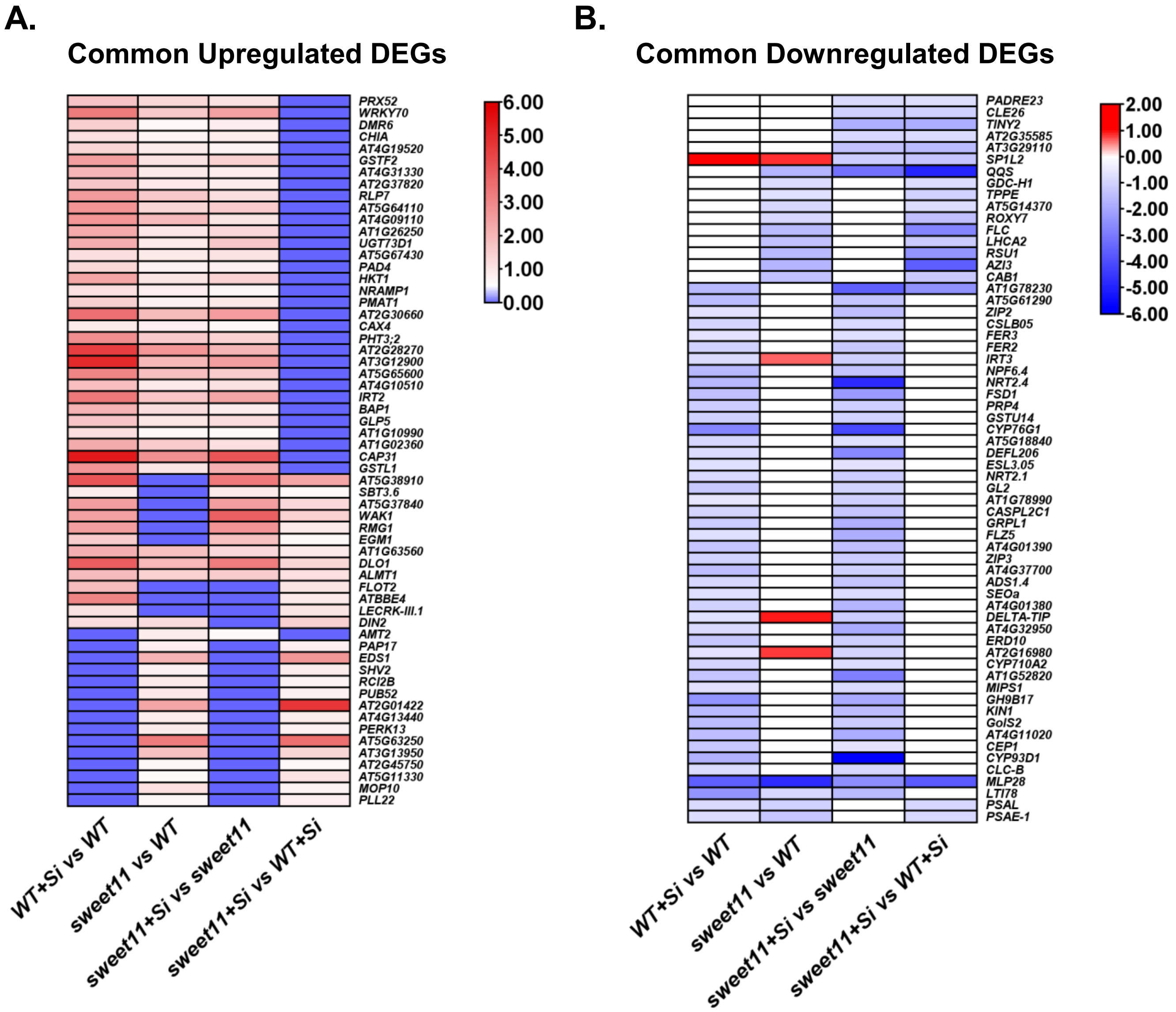


**Fig. S11. Expression of common DEGs in *S. indica*-colonized WT and *sweet11* mutant roots. A. Common upregulated DEGs:** Heat map was prepared using Log2FC at cutoff of ± 0.58. All DEGs in *S. indica*-colonized *sweet11* roots, *S. indica*-colonized WT roots and *sweet11* control were as input in TB tool. **B. Common downregulated DEGs:** Heat map was prepared using Log2FC at cutoff of ± 0.58. All DEGs in *S. indica*-colonized *sweet11* roots, *S. indica*-colonized WT roots and *sweet11* control were as input in TB tool.

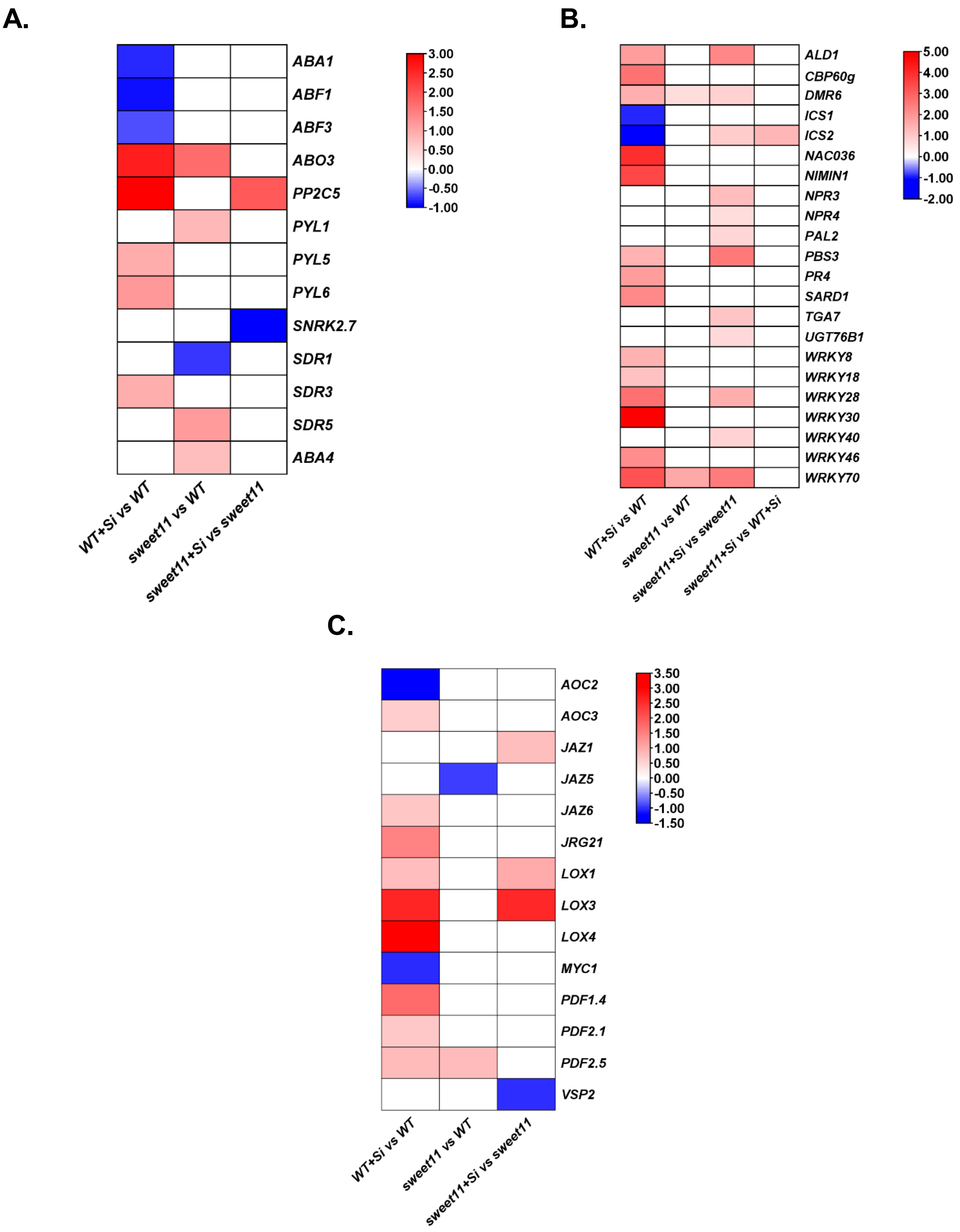
A.
ABA1
ABF1
ABF3
ABO3
PP2C5
PYL1
PYL5
PYL6
SNRK2.7
SDR1
SDR3
SDR5
ABA4
WT+Si vs WT
sweet11 vs WT
sweet11+Si vs sweet11
3.00
2.50
2.00
1.50
1.00
0.50
0.00
-0.50
-1.00
B.
ALD1
CBP60g
DMR6
ICS1
ICS2
NAC036
NIMIN1
NPR3
NPR4
PAL2
PBS3
PR4
SARD1
TGA7
UGT76B1
WRKY8
WRKY18
WRKY28
WRKY30
WRKY40
WRKY46
WRKY70
WT+Si vs WT
sweet11 vs WT
sweet11+Si vs sweet11
sweet11+Si vs WT+Si
5.00
4.00
3.00
2.00
1.00
0.00
-1.00
-2.00
C.
AOC2
AOC3
JAZ1
JAZ5
JAZ6
JRG21
LOX1
LOX3
LOX4
MYC1
PDF1.4
PDF2.1
PDF2.5
VSP2
WT+Si vs WT
sweet11 vs WT
sweet11+Si vs sweet11
3.50
3.00
2.50
2.00
1.50
1.00
0.50
0.00
-0.50
-1.00
-1.50

**Fig. S12. Transcripts level of ABA and SA related genes in Arabidopsis wild-type (WT) and *sweet11* mutant 7 dpi in roots upon *S. indica* colonization. A.** Heat map of ABA biosynthesis and ABA signaling related genes. **B.** Heat map of SA biosynthesis and ABA signaling genes. **C.** Heat map of JA biosynthesis and JA signaling genes. Differential expression was calculated over control and the log2fold-change change was taken at cutoff of ±0.58.

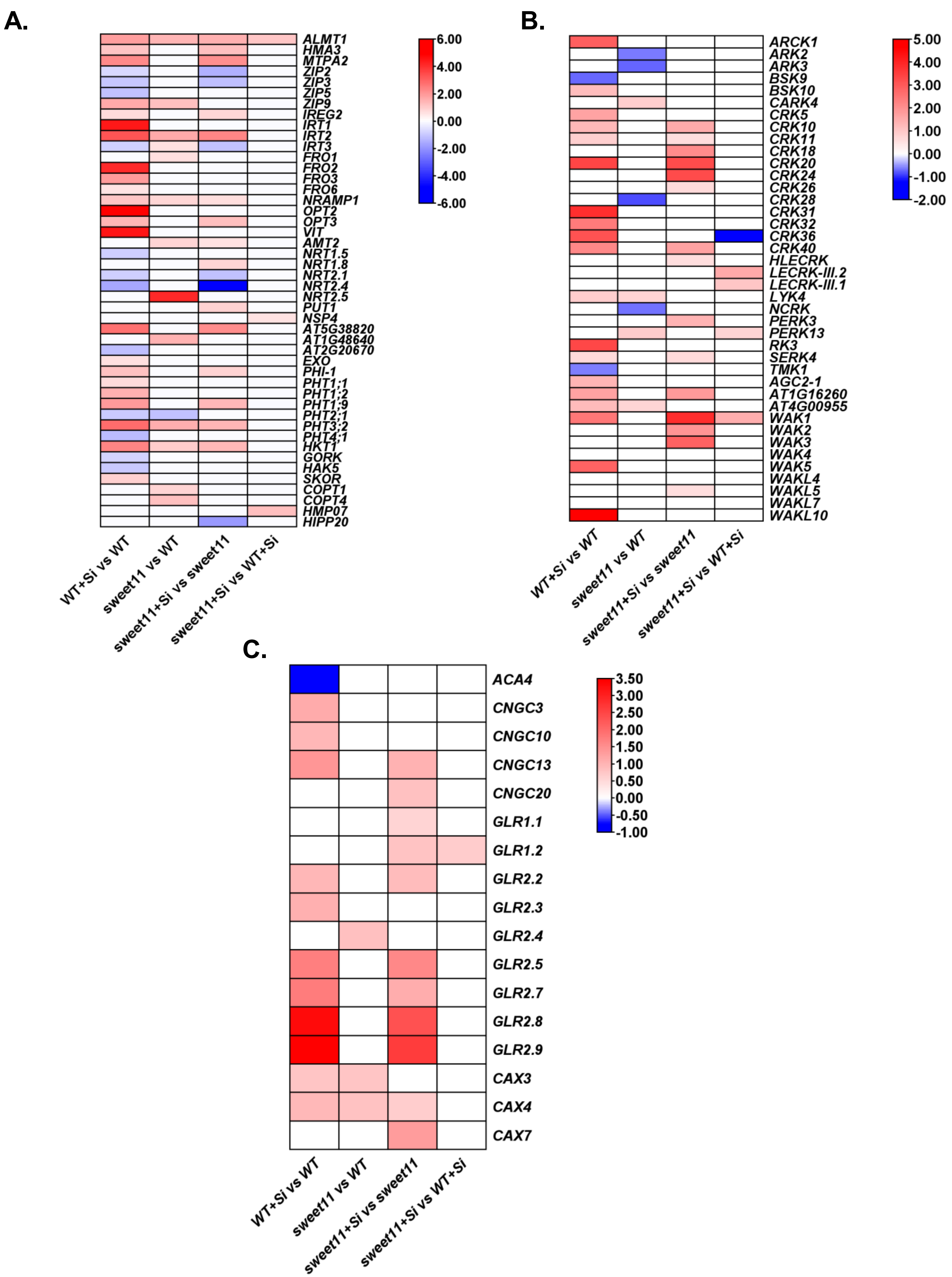
A.
ALMT1
HMA3
MTPA2
ZIP2
ZIP3
ZIP5
ZIP9
IREG2
IRT1
IRT2
IRT3
FRO1
FRO2
FRO3
FRO6
NRAMP1
OPT2
OPT3
VIT
AMT2
NRT1.5
NRT1.8
NRT2.1
NRT2.4
NRT2.5
PUT1
NSP4
AT5G38820
AT1G48640
AT2G20670
EXO
PHI-1
PHT1;1
PHT1;2
PHT1;9
PHT2;1
PHT3;2
PHT4;1
HKT1
GORK
HAK5
SKOR
COPT1
COPT4
HMP07
HIPP20
6.00
4.00
2.00
0.00
-2.00
-4.00
-6.00
WT+Si vs WT
sweet11 vs WT
sweet11+Si vs sweet11
sweet11+Si vs WT+Si
B.
ARCK1
ARK2
ARK3
BSK9
BSK10
CARK4
CRK5
CRK10
CRK11
CRK18
CRK20
CRK24
CRK26
CRK28
CRK31
CRK32
CRK36
CRK40
HLECRK
LECRK-III.2
LECRK-III.1
LYK4
NCRK
PERK3
PERK13
RK3
SERK4
TMK1
AGC2-1
AT1G16260
AT4G00955
WAK1
WAK2
WAK3
WAK4
WAK5
WAKL4
WAKL5
WAKL7
WAKL10
5.00
4.00
3.00
2.00
1.00
0.00
-1.00
-2.00
WT+Si vs WT
sweet11 vs WT
sweet11+Si vs sweet11
sweet11+Si vs WT+Si
C.
ACA4
CNGC3
CNGC10
CNGC13
CNGC20
GLR1.1
GLR1.2
GLR2.2
GLR2.3
GLR2.4
GLR2.5
GLR2.7
GLR2.8
GLR2.9
CAX3
CAX4
CAX7
3.50
3.00
2.50
2.00
1.50
1.00
0.50
0.00
-0.50
-1.00
WT+Si vs WT
sweet11 vs WT
sweet11+Si vs sweet11
sweet11+Si vs WT+Si

**Fig. S13. Transcripts level of nutrient transport, receptors and calcium channels related genes in Arabidopsis wild-type (WT) and *sweet11* mutant 7 dpi in roots upon *S. indica* colonization. A.** Heat map of nutrient transport related genes. **B.** Heat map of receptors and cell membrane associated kinases related genes. **C.** Heat map of calcium channels DEGs. Differential expression was calculated over control and the log2fold-change change was taken at cutoff of ±0.58
